# A neural geometry theory comprehensively explains apparently conflicting models of visual perceptual learning

**DOI:** 10.1101/2023.11.13.566963

**Authors:** Yu-Ang Cheng, Mehdi Sanayei, Xing Chen, Ke Jia, Sheng Li, Fang Fang, Takeo Watanabe, Alexander Thiele, Ru-Yuan Zhang

## Abstract

Visual perceptual learning (VPL), defined as long-term improvement in a visual task, is considered a crucial tool for elucidating underlying visual and brain plasticity. However, the identification of a unified theory of VPL has long been controversial. Multiple existing models have proposed diverse mechanisms, including improved signal-to-noise ratio, changes in tuning curves, and reduction of noise correlations, as major contributors to improved neural representations associated with VPL. However, each model only accounts for specific aspects of the empirical findings, and there exists no theory that can comprehensively explain all empirical results. Here, we argue that all neural changes at single units can be conceptualized as geometric transformations of population response manifolds in a high-dimensional neural space. This approach enables conflicting major models of VPL to be quantitatively tested and compared within a unified computational theory. Following this approach, we found that changes in tuning curves and noise correlations, as emphasized by previous models, make no significant contributions to improved population representations by visual training. Instead, we identified neural manifold shrinkage due to reduced trial-by-trial neural response variability, a previously unexplored factor, as the primary mechanism underlying improved population representations. Furthermore, we showed that manifold shrinkage successfully accounts for learning effects across various domains, including artificial neural responses in deep neural networks trained on typical VPL tasks, multivariate BOLD signals in humans, and multi-unit activities in monkeys. These converging results suggest that our neural geometry theory offers a quantitative and comprehensive approach to explain a wide range of empirical results and to reconcile previously conflicting models of VPL.

## INTRODUCTION

Adapting to new visual environments is crucial for an organism’s survival in its environment. This ability is well exemplified by visual perceptual learning (VPL), which is defined as long-term performance enhancements resulting from visual experience 1,2. Therefore, elucidating the mechanisms underlying VPL may contribute to an understanding of how organisms adapt to new environments. However, despite years of research in systems neuroscience, psychophysics and machine learning, the mechanisms behind VPL remain mysterious.

It is widely acknowledged that visual training enhances behavioral performance and refines neural representations in neural populations. Previous studies using human neuroimaging and monkey neurophysiology have demonstrated a significant improvement in the fidelity of stimulus encoding within population responses 3-5. These findings strongly support the theory that enhanced signal-to-noise ratios serve as a potent computational mechanism for refined neural representations associated with VPL (Fig. 1E) 6-8. However, the precise mechanisms underlying how visual training refines neural representations remain elusive. Several conflicting models have been proposed based on neural changes associated with VPL. One model suggests that VPL is associated with changes in population representations resulting from changes in neuronal tuning curves, as evidenced by sharpened orientation tuning curves in monkey visual cortex 9,10. Another model assumes that changes in population representations result from a reduction in trial-by-trial co-variation of neuronal firing rate, known as noise correlations, which have been observed in association with VPL in both monkeys and songbirds 11-14.

**Figure 1.**
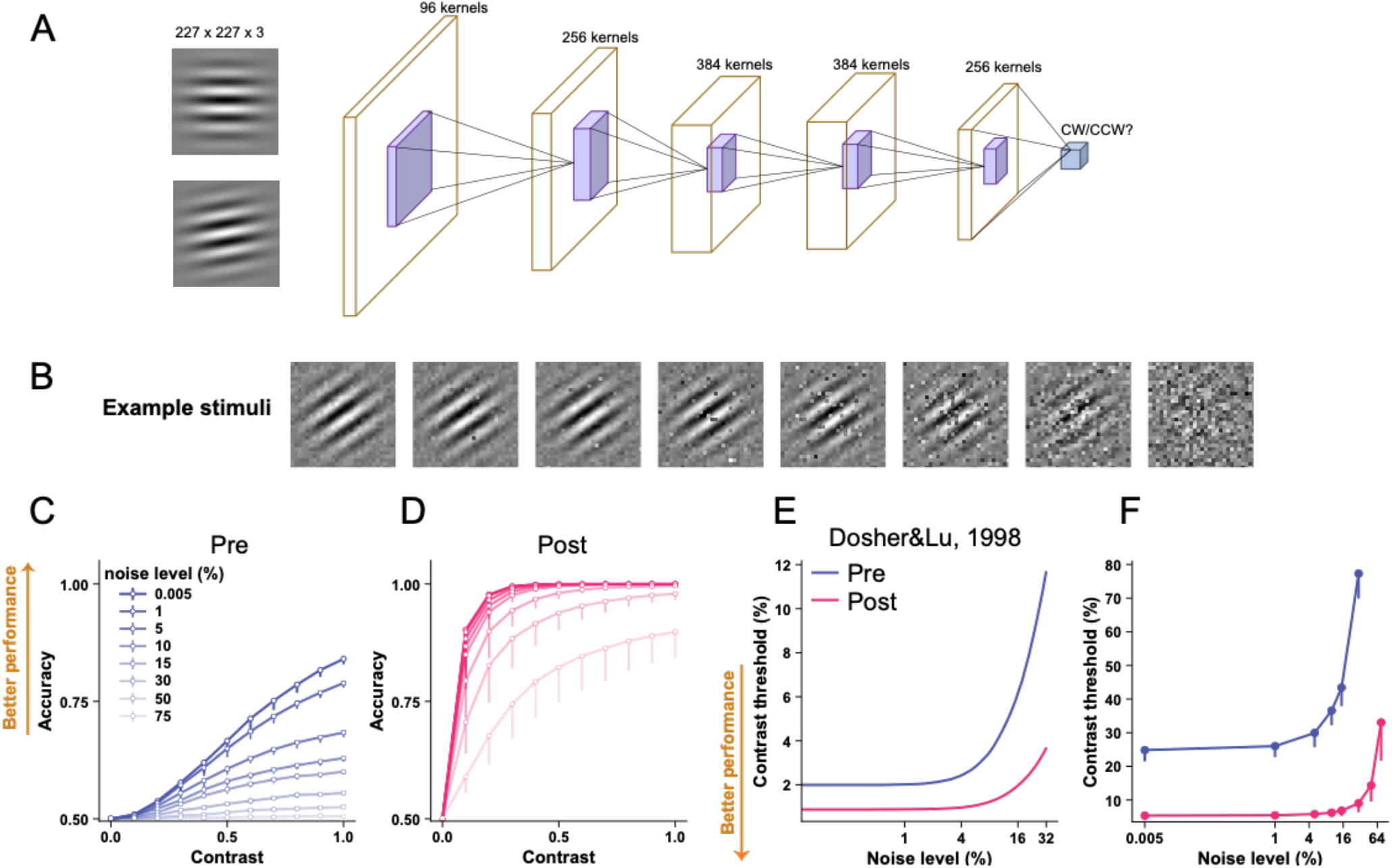
A DCNN (***A***) is trained on an orientation discrimination task with Gabor stimuli embedded in different levels of image noise (***B***). Orientation discrimination accuracy is significantly improved from pre-(***C***) to post-test (***D***). Training induces a downshift of the Threshold vs. Noise function (***F***), an effect that is qualitatively similar to existing human psychophysical results (***E***, corresponds to the 70.7% accuracy condition in Fig. 1 of ref. 7). The absolute quantitative differences between ***E*** and ***F*** may be due to differences in the overall signal-to-noise ratio or the number of layers and units between the human visual system and the DCNN.

The primary conceptual gap in reconciling the conflicting models of VPL lies in their focus on mechanisms proposed at the single-unit level (e.g., changes in tuning curves or noise correlations), whereas the effects of these mechanisms in VPL should be evaluated at the population level (i.e., improved population representations). Although VPL is indeed associated with changes in both single-unit responses and improved population representations, it remains controversial whether changes in single-unit responses are the actual *cause* of improved population representations. It is possible that changes in single-unit responses associated with VPL are merely by-products of underlying processes. While these conflicting models of VPL capture certain aspects of the empirical findings, they fail to generate falsifiable predictions about how changes in single-unit responses contribute to improved population representations.

A significant obstacle to comparing the conflicting models of VPL is the complex interactions between different aspects of single-unit responses (e.g., tuning curves and noise correlations) on population representations. Computational neuroscience research has elucidated that the impact of noise correlations on population representations heavily depends on its interaction with tuning curves 15. It is important to note that reduced noise correlations do not inherently enhance information in a neural population 16-18. Moreover, the challenge is exacerbated by the fact that their interaction effects are even changing rather than remaining stable throughout a training process, because both noise correlations and tuning curves change. These dynamic changes further complicate the understanding of how training affects their interactions. To overcome this, a comprehensive computational theory is imperative to quantify and disentangle the effects of different changes in single-unit responses, such as sharpened tuning curves and reduced noise correlations, on neural representations at the population level.

To comprehensively explain these conflicting models, we developed a neural geometry theory of VPL. In this theory, trial-by-trial population responses elicited by two stimuli for discrimination form two differentiable manifolds in a high-dimensional neural space. In this space, changes in single-unit responses (e.g., tuning curves, Fano factor, and noise correlations) can be interpreted as changes in several fundamental and measurable geometric properties (e.g., centroids, size, orientations) of neural manifolds. This approach allows quantitative comparisons of conflicting models of VPL and assessments of their contributions to population representations within the same computational framework. Thus, this theory directly bridges single-unit responses and population representations, and offers a normative account of the potential neural mechanisms underlying VPL. Specifically, this theory proposes four possible training-induced geometric changes (signal enhancement, manifold shrinkage, signal rotation, and manifold warping) that can summarize all previous models of VPL. Thus, improved population representations can be achieved by one or a combination of the four interpretable mechanisms.

To assess this neural geometry theory, we trained deep convolutional neural network (DCNN) on two typical VPL tasks—orientation discrimination and motion direction discrimination. First, the DCNNs successfully replicated a wide range of psychophysical and imaging findings in humans, as well as neurophysiological findings in monkeys. Second, analyses of the geometric mechanisms mentioned above suggest that changes in both tuning curves and noise correlations are indeed present in VPL. Third, and most importantly, our analysis further revealed that neither changes in tuning curves nor changes in noise correlations at the single-unit level contributed significantly to improved population representations, despite their strong association with VPL. Surprisingly, we found that neural manifold shrinkage induced by reduced response variability emerged as the primary mechanism driving VPL. We conducted additional tests to examine the predictions derived from the geometry theory across different levels of measurement in different spaces. Remarkably, we found that the geometry theory incorporating manifold shrinkage aligned closely with BOLD response changes associated with VPL of motion direction discrimination in humans, as well as the electrophysiological population response changes associated with VPL of contrast discrimination in monkey V4.

## RESULTS

### Orientation discrimination learning improves behavioral performance of DCNN

To elucidate the neurocomputational mechanisms of VPL, we trained a DCNN (Fig. 1A) to perform a classical orientation discrimination task 7. Similar to the neural network in ref. 19, this neural network inherits the first five convolutional layers of AlexNet, which was pre-trained on ImageNet 20. To emulate the decision stage of orientation discrimination, we added a linear decoding layer and used the logistic function to classify the activity of the decision unit into a binary perceptual choice (i.e., clockwise or counterclockwise rotation of the target stimulus relative to the reference stimulus). Importantly, similar to previous psychophysical studies 7,21, we systematically manipulated the level of input image noise (Fig. 1B). The network was trained on stimuli with multiple noise and contrast levels (see Methods and Materials for training details). To evaluate the performance of the neural network and compare it directly with human psychophysical results, we assessed orientation discrimination accuracy as a function of stimulus contrast and noise (i.e., behavioral psychometric functions in Fig. 1C&D), and further derived contrast thresholds as a function of image noise level (Fig. 1F, Threshold vs. Noise, TvN function). We found that training significantly improved the network performance in this task in almost all stimulus contrast and noise conditions. Importantly, training uniformly reduced contrast thresholds at all noise levels (Fig. 1F). This downshift of TvN functions is consistent with the well-established psychophysical result (replotted in Fig. 1E) as previously observed in humans 7,8.

### Orientation discrimination learning refines neural population representations

Having established the behavioral signature of VPL in our DCNN, we next sought to understand the effects of visual training on population representations in the network. We simulated many trials elicited by the two orientation stimuli for fine discrimination and performed multivariate decoding analyses in each layer. Training improved decoding accuracy in later layers (layers 3-5, Fig. 2F). The learning effect is more pronounced in later layers possibly because the benefits of visual training accumulate from early to later layers. More formally, we calculated linear Fisher information, a classical metric in computational neuroscience, to quantify stimulus information (i.e., how well the two stimuli can be discriminated based on population responses, see Methods and Materials) in each layer before and after training. The amount of sensory information represented in later layers was indeed enhanced by training (Fig. 2G). Such refined neural representation at the population level is consistent with the decoding results based on both macroscopic cortical activity in humans 3,4,22 and multi-unit spiking activity in monkeys 13,14.

**Figure 2.**
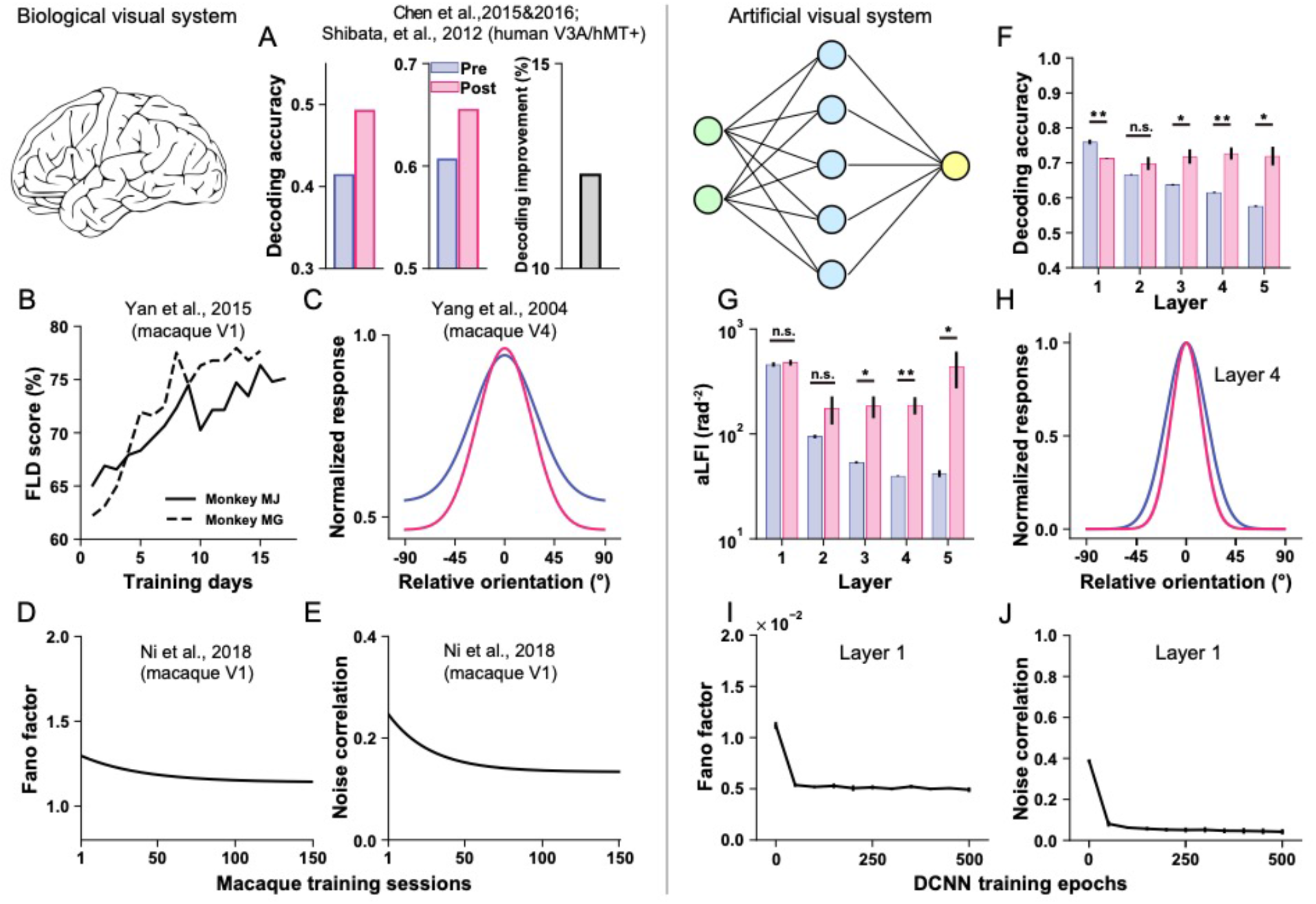
Neural correlates of VPL in humans (***A***), monkeys (***B*-*E***), and in our DCNN (***F*-*J***). Visual training improves stimulus decoding accuracy in related regions in the human brain (***A***) and decoding scores of Fisher’s linear discriminant (FLD) in monkey V1 (***B***). Visual training sharpens orientation tuning curves of neurons in monkey V4 (***C***), and also reduces Fano factors and interneuron noise correlations (***D&E***). Similar results are observed in the DCNN: network training also improves decoding accuracy in layers 3-5 (***F***), and averaged linear Fisher information (aLFI, total information in each layer divided by the number of units in that layer) (***G***) in layers 3-5. Training sharpens orientation tuning curves of units in layers 1-4 in the DCNN (results of layer 4 only are shown in ***H***). Similar reduction of Fano factors and noise correlations are observed in the DCNN (results of layer 1 only are shown in ***I&J***). The data shown in ***H****-****J*** are the median value across units in a layer. The results of all five layers are shown in Supplementary Fig. S1. Panels ***A-E*** are reproduced by the data points shown in the original papers. Error bars and error shadings in panels ***F-J*** represent the S.E.M. across four reference orientations (error shadings in ***H*** are small and barely visible).

### Orientation discrimination learning changes response properties of individual units

In addition to characterizing VPL on the population level, we sought to investigate the neural correlates at the single-unit level.

Importantly, we found that three key neural signatures of VPL as documented in the neurophysiological literature emerge naturally from the neural network training. First, training modestly sharpened the tuning curves of artificial neurons in layers 1 to 4 (Fig. 2H& and Supplementary Fig. S1). The finding of changed tuning curves has been reported previously in refs. 9,10,23 (Fig. 2C, but see also null results in ref. 24). Second, we observed a significant decrease in Fano factor of individual units in all five layers (Fig. 2I and Supplementary Fig. S1), a phenomenon indicating an increased signal-to-noise ratio of individual neuronal responses. This has been reported in both humans 25 and monkeys 11,23 (Fig. 2D). The sharpened tuning curve and reduced Fano factor are also consistent with theoretical modeling 19. Third, training reduced trial-by-trial noise correlations between units in all five layers (Fig. 2J and Supplementary Fig. S1), a finding also consistent with several empirical results in monkeys 11-14. Critically, we also found that the reduction in noise correlation depended on tuning similarity. Learning reduced the noise correlations between units with similar tunings (i.e., positive signal correlations) and increased the noise correlations between units with opposite tunings (i.e., negative signal correlations) (Supplementary Fig. S2). Previous theoretical work has suggested that the former type of noise correlations is detrimental for information coding and the latter type is beneficial 15,16. The pattern of reduced detrimental and increased beneficial noise correlations has been discovered with learning tasks in songbirds 26 and with attention tasks in monkeys 27.

In addition to these classical neurophysiological findings in VPL, our network also captures some important response properties of sensory neurons in the primate early visual system. First, the relationship between the Fano factor and orientation tuning of the artificial neurons bears strong resemblances to the empirical measures of V1 neurons in monkeys 28 (Supplementary Fig. S2). Second, we found a positive relationship between signal correlation and noise correlation among artificial neurons in all layers (Supplementary Fig. S2). This relationship has also recently been documented as a ubiquitous phenomenon in both electrophysiological 29-31 and human imaging 17,18,32 studies.

Taken together, our neural network reproduces a wide range of psychophysical, human imaging, and monkey neurophysiological findings in VPL. Although these similarities are qualitative and not quantitative, these results suggest that our DCNN allows us to explore neurocomputational mechanisms that may be difficult to elucidate in empirical experiments.

### Neural correlates of four computational mechanisms under the neural geometry theory of VPL

How would improved sensory discrimination manifest in high-dimensional population responses? In the simplified one-dimensional scenario (Fig. 3A), the classical signal detection theory posits that better sensory discrimination can be achieved by either increasing the distance between the means (i.e., signal enhancement) and/or decreasing the variance (i.e., noise reduction) of the two response distributions, because the former only changes the tuning (i.e., mean) of a response unit and the latter only reduces the variance around a fixed mean. In multivariate population responses, the two stimuli to be discriminated instead generate two multivariate response distributions (i.e., neural manifold) in a high-dimensional neural space whose dimension corresponds to the dimension of a population (Fig. 3B&C). In a simplified visualization in 2D space (i.e., assuming a population of only two units, Fig. 3D), the two distributions are elliptical due to noise correlations between units. We refer to the vector connecting the mean of the two distributions as the *signal vector* and its modulus length (i.e., the Euclidean distance between the two manifold centroids) as the *signal separation*.

**Figure 3.**
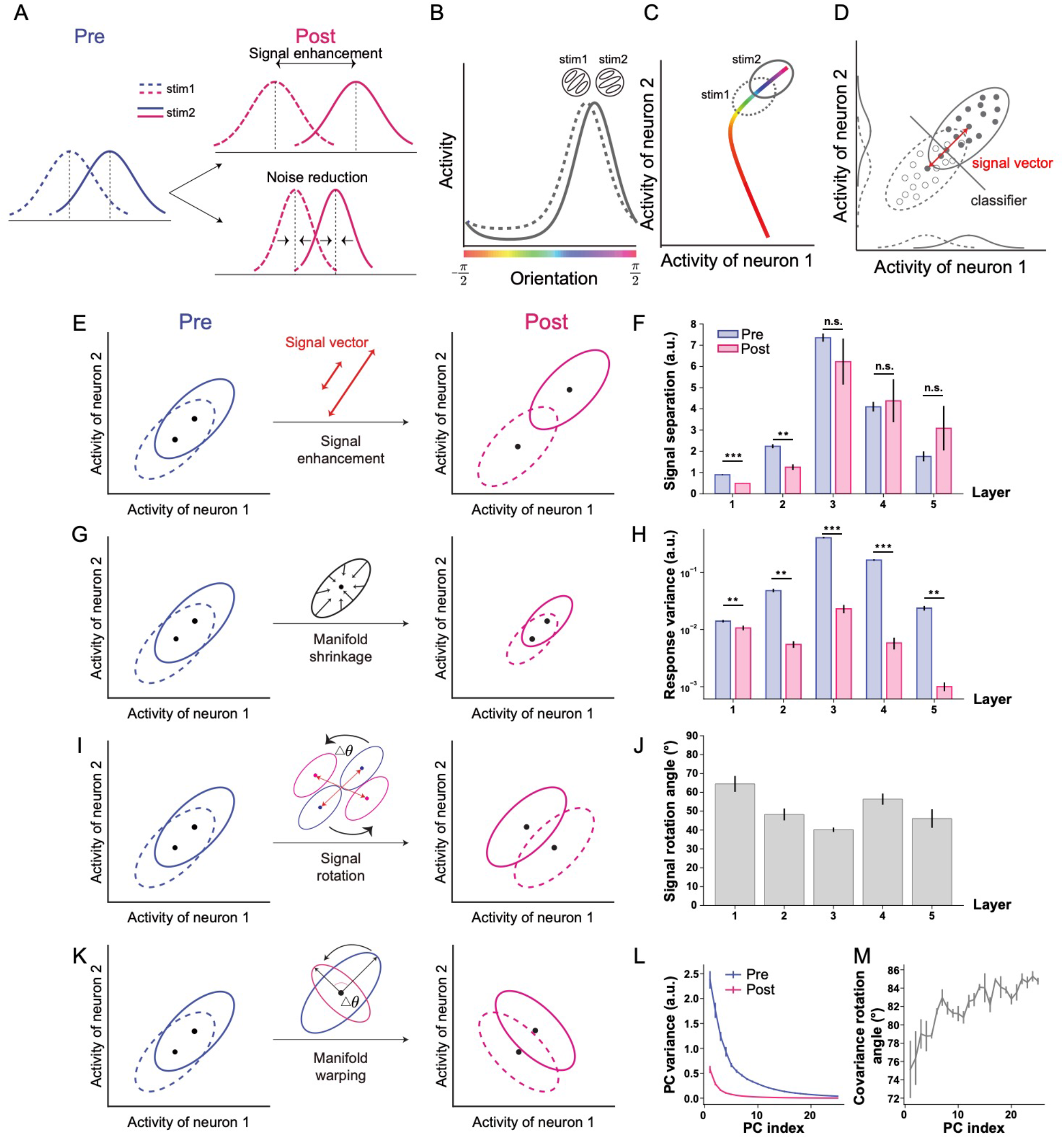
Four possible mechanisms of VPL in neural populations. ***A***. According to classical signal detection theory, better sensory discriminability can be represented as a larger separation between two response distributions. Signal enhancement predicts enlarged distances between the mean of the two distributions while noise reduction predicts reduced variance between the two response distributions. ***B***. Stimulus orientation as a continuous stimulus variable can evoke population responses with the same shape as individual tuning curves. ***C***. If we continuously sweep the orientation value, the mean of the population responses forms a manifold in a high-dimensional neural space with the number of dimensions being equal to the number of units. The mean population responses to the two stimuli in a discrimination task are two points on the manifold. ***D***. In realistic population responses, the trial-by-trial population responses to the two stimuli form two high-dimensional response distributions (i.e., neural manifolds). The manifolds look elliptical rather than spherical due to pairwise noise correlations between units. In this high-dimensional neural space, the signal enhancement mechanism predicts an increased Euclidean distance (i.e., signal separation, ***E***) between two high-dimensional response distributions. However, no significant increase in signal separation is observed in any of the five layers (signal separation decreases in the first two layers, ***F***). The manifold shrinkage mechanism predicts reduced variance (***G***) of the two neural manifolds. This is observed in all five layers (***H***). ***I***. The signal rotation mechanism predicts that the positions of the centroid (i.e., mean) of the two manifolds are changed by training. ***J***. The rotation angle ranges from 50º∼70º in all five layers, supporting the existence of signal rotation. ***K***. The manifold warping mechanism predicts that training changes the shape of the population response covariance. ***L***. Indeed, training mostly reduces the variance of the high-variance principal components of the population responses. The principal components (PC, only show PCs that account for >99% of the total variance) are ranked from high to low variance. ***M***. More importantly, the directions of the principal components rotate significantly from pre-to post-test.

In the high-dimensional neural space, our neural geometry theory of VPL proposes that visual training improves sensory discrimination by shaping some fundamental geometric properties of the neural manifolds. Here, under this theory, there exist only four possible mechanisms to separate two neural manifolds (Eq. 4 in Methods and Materials). First, according to the classical signal detection theory, the *signal enhancement* mechanism predicts an increased Euclidean distance between the centroids of the two neural manifolds (Fig. 3E). However, we found that the signal separation between the two manifolds did not significantly increase with learning in all five layers, and even slightly decreased in the first two layers (Fig. 3F). Second, the *manifold shrinkage* mechanism predicts that visual training reduces the trial-by-trial response variance of units, thereby reducing the size of the manifolds (Fig. 3G). This is what we found in all five layers (Fig. 3H). These results argue that manifold shrinkage plays a more important role than signal enhancement in VPL. We further included two novel mechanisms that can only occur in high-dimensional neural space and lead to increased discriminability of two response distributions. In the third mechanism, although visual training did not increase signal separation, it may change the positions of the centroids of the two manifolds and consequently increase discriminability due to the elliptical shape of the manifolds (Fig. 3I). Interestingly, we found that the signal vectors in each layer were rotated by ∼50°-70° after training (Fig. 3J). We call this mechanism *signal rotation*. Fourth, visual training can warp the shapes of the high-dimensional neural manifolds while keeping the size of the manifolds unchanged. As evidenced by the change of covariance structure, we found that visual training systematically warped the shape (i.e., covariance structures) of the high-dimensional neural manifolds (Fig. 3K-M). We refer to this mechanism as *manifold warping*. Note that manifold warping includes both the changes in correlation structures and the redistribution of variances across individual units, while holding the total variance constant. It is manifold shrinkage that attenuates the total variance.

In summary, our neural geometry theory of VPL summarizes possible mechanisms of improved sensory discrimination into explicit and interpretable changes of geometric properties in high-dimensional neural space. We present four possible computational mechanisms predicted by the neural geometry theory of VPL and identify neural evidence for three of them (i.e., except signal enhancement) in our DCNN training. Note that these four mechanisms (or a combination thereof) encompass all possible scenarios by which learning can potentially improve sensory discrimination.

### Information-theoretic analyses reveal the key role of manifold shrinkage in orientation discrimination learning

Given the four possible mechanisms (i.e., signal enhancement, manifold shrinkage, signal rotation, and manifold warping) and their complex interaction effects, how can we delineate their respective contributions to improved population representations? Here, we use linear Fisher information to quantify manifold separability. Besides, we introduce a stepwise approach to further disentangle the respective contributions of the four possible mechanisms. Specially, their respective contributions are assessed by sequentially allowing only one mechanism to occur and quantifying its endowed changes in the linear Fisher information of whole populations (Fig. 4A). For example, as shown in Fig. 4, we first calculate how much information is enhanced by considering only the signal enhancement scenario, then by considering both signal enhancement and manifold shrinkage, and so on until all four mechanisms are included. Here sensory information refers to manifold separability. We emphasize that this stepwise approach is intended only to facilitate the understanding of the computational underpinning of each mechanism. It does not imply that these mechanisms must occur sequentially over the course of visual training.

**Figure 4.**
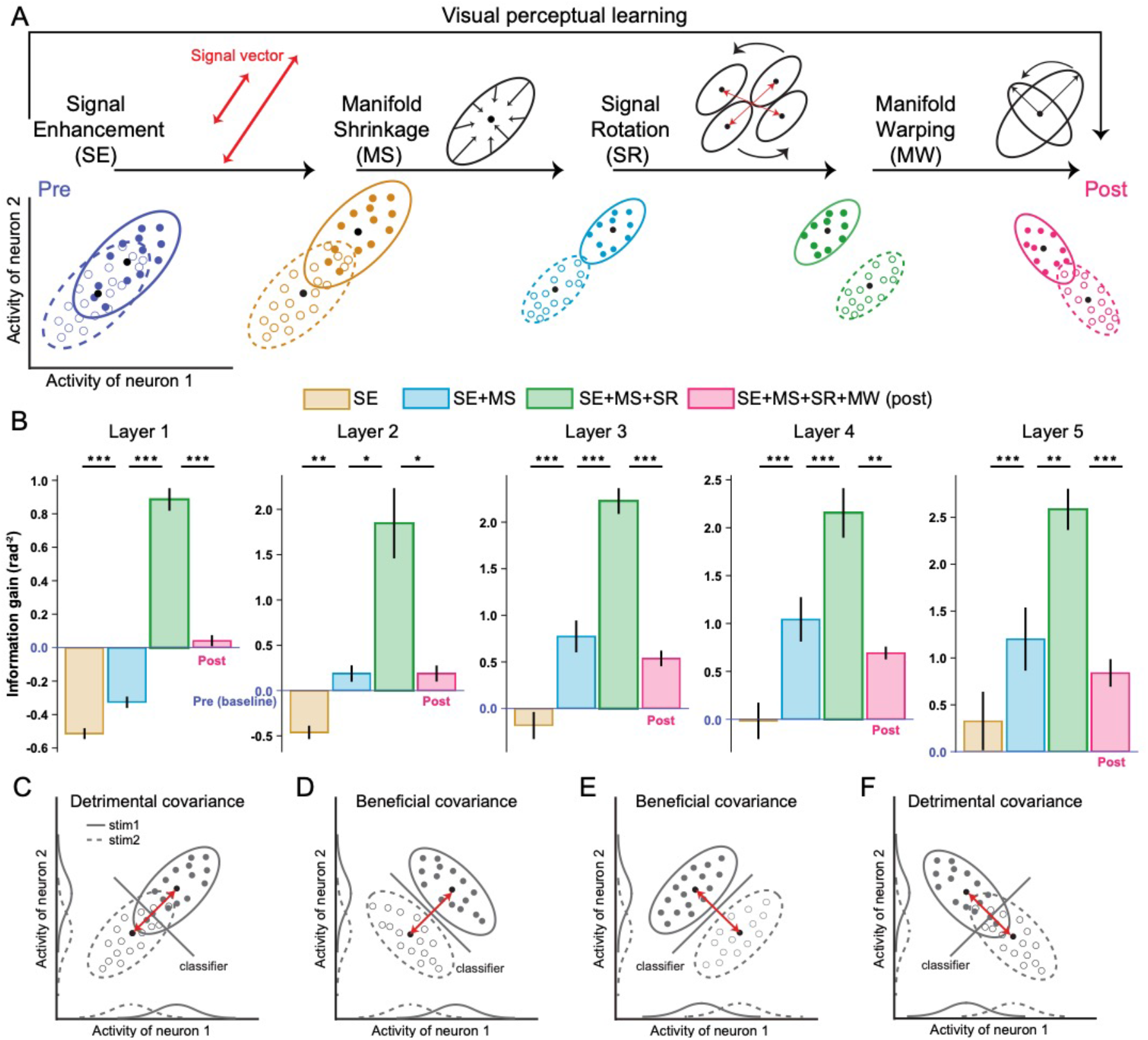
***A***. The effects of four mechanisms on population representations are decomposed into four distinct steps (***A***). ***B*** shows the effects on information gain by sequentially adding each of the four mechanisms in each layer. For example, the increase in height from the brown to the blue bars indicates the positive contribution of manifold shrinkage to encoded stimulus information. ***C-F*** illustrate the strong interaction effects between covariance and signal vector. For distributions with identical covariance (***C*** and ***E***; ***D*** and ***F***), detrimental (***C*** or ***F***) or beneficial (***D*** or ***E***) effects on discriminability are possible, depending the signal vector. Similarly, the effects of the signal vector also depend on its relative geometry to the axis of covariance.

Interestingly, not all mechanisms have a positive effect on population representations. We found that the effect of signal enhancement is minimal in all five layers. This mechanism even reduces stimulus information in layers 1&2. This is consistent with the reduced Euclidean distance in the first two layers (Fig. 3F). Manifold shrinkage significantly enhances stimulus information in almost all layers. This makes sense because the decreased variance reduces the overlap between the two manifolds. Interestingly, we found that signal rotation appears to enhance stimulus information (Fig. 4B, green bars). This is because rotation of the signal vectors disrupts their relative parallelism to the covariance direction at pre-test, making them more orthogonal. Such changes increase the apparent information. However, the effect becomes minimal when manifold warping is further considered (Fig. 4B, magenta bars) because visual training also warps the covariance direction to realign it with the post-test signal vector, thereby reducing stimulus information.

We want to highlight two important aspects of such stepwise analyses. First, we use signal rotation and manifold warping to reflect changes in the direction of the signal vector and the shape of the covariance, which are theoretically independent (Fig. 3I&K). But what matters for the fidelity of population representations is the relative geometric relationship between covariance and signal vector rather than their absolute magnitude 15,16 (Fig. 4C-F). Given that learning changes both signal vector and covariance, we argue that their contributions to improved population representations should be evaluated together. In Fig. 4B, signal rotation and manifold warping appear to have strong positive and negative effects on information, respectively. However, their joint effect is modest (i.e., the magenta bars compared to the blue bars in Fig. 4B). Thus, it can be argued that manifold shrinkage is the primary contributor to improved stimulus information (also see Discussion).

Second, the results are not affected by the order in which the mechanisms are applied changes. As mentioned above, the effects of signal rotation and manifold warping should be evaluated together, but the effects of signal enhancement and manifold shrinkage are independent. Hence, changing the order does not affect our results as long as signal rotation and manifold warping are carried out consecutively. In fact, the apparent positive and negative effects of signal rotation and manifold warping in the stepwise analyses can be reversed when their order is switched, but both orders actually reflect identical underlying changes (see detailed analyses in Supplementary Fig. S6).

Taken together, we propose an interpretable and quantitative neural geometry theory of VPL where visual training refines the geometry of representations in a high-dimensional neural space. Specially, our neural geometry theory predicts four different computational mechanisms of VPL. Our neural network allows us to readily measure the activity of all our simulated units across the visual hierarchy. Using the neural network, we first verified the existence of three of four possible mechanisms, and we performed systematic information-theoretic analyses to disentangle the effects of them. Most importantly, we found that manifold shrinkage in population responses was the key mechanism underlying the improved population representations induced by visual training in the DCNN.

### Neural geometric changes of motion direction discrimination learning in DCNN

The above analyses focus only on a classical VPL task—orientation discrimination and a specific neural network structure—a six-layer convolutional neural network. In this section, we switch to motion VPL—another sensory domain that is also widely used in psychophysical 33,34, human imaging 3,4, and neurophysiological studies 35. Importantly, motion VPL involves the processing of both spatial and temporal signals rather than merely static spatial information in orientation learning. Similarly, we inherited the first six layers of the pre-trained C3D network 36 and trained the neural network to perform a motion direction discrimination task commonly used in psychophysics (see Methods and Materials for stimulus and training details).

In the motion DCNN, we found similar mechanisms as in the orientation discrimination learning task. First, motion direction discrimination training significantly improved the behavioral performance of the network (Fig. 5B). Second, training also enhanced decoding accuracy and averaged linear Fisher information in later layers (Fig. 5C&D), suggesting that such training refines stimulus representation at the population level. Third, the effects of motion direction discrimination training on individual units in layer 6 are also pronounced (see results for all six layers in Supplementary Fig. S3). We found that training reduced Fano factor (Fig. 5E) and noise correlations (Fig. 5F). Fourth, training did not significantly improve signal separation (Fig. 5G) but markedly reduced response variance (Fig. 5H). In addition, motion direction discrimination training also induced two novel mechanisms—signal rotation (Fig. 5I) and manifold warping (Fig. 5J&K). Most importantly, the four mechanisms induced by the training had similar respective contributions to population representations (Fig. 5L).

**Figure 5.**
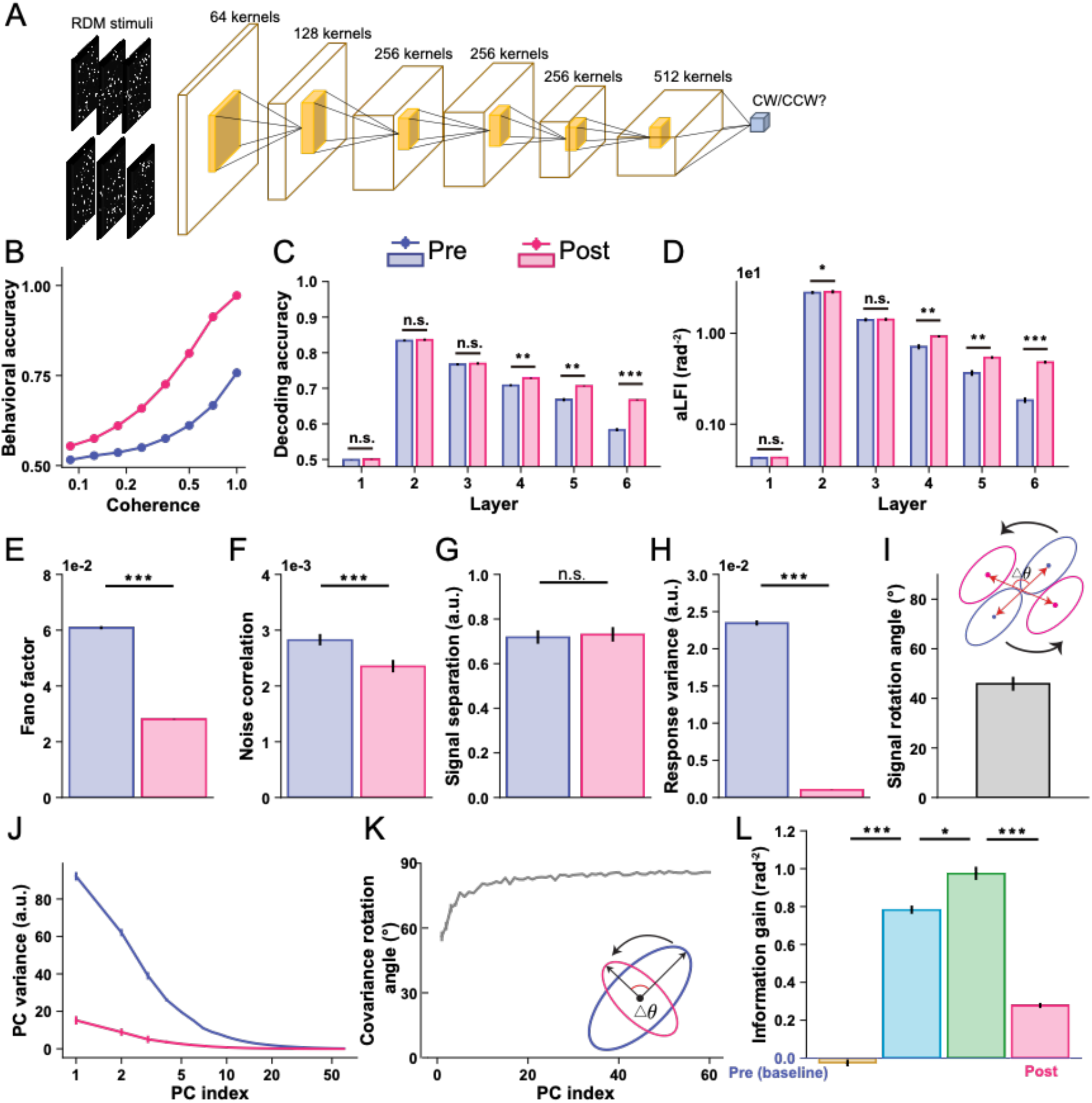
DCNN modeling of motion VPL. The DCNN (***A***) of motion VPL uses 3D convolutions to process video stimuli. Here, we simplify the 4D feature maps in each convolutional layer and show them as 3D maps only for illustration purposes. Training significantly improves DCNN direction discrimination performance (***B***), decoding accuracy (***C***), and averaged linear Fisher information (***D***). For single-unit analyses, motion direction discrimination training also reduces the Fano factor (***E***) and noise correlation (***F***) in layer 6. Similar to orientation discrimination training, motion direction discrimination training does not significantly enhance signal separation (***G***), but significantly rotates the position of the two distributions in layer 6 (***I***). Importantly, training clearly reduces the response variance in layer 6 (***H***). Specifically, training reduces the variance of the high variance PCs and rotates the directions of all PCs, indicating a significant effect of manifold warping in layer 6 (***J&K***). The pattern of information gain associated with the four possible mechanisms (***L***) is consistent with that of orientation discrimination training. See results for all six layers in Supplementary Fig. S3. Note that some error bars are very small and barely visible.

### Neural geometric changes of motion direction discrimination learning in the human brain

Above, our DCNNs help verify, quantify and disentangle the four possible mechanisms proposed by our neural geometry theory. The converging results in the DCNNs of orientation and motion direction discrimination, and the remarkable agreement between our DCNNs and existing empirical neuroscientific findings support the biological plausibility of our DCNNs. However, it remains unknown whether these predictions are only present in the DCNNs and have no biological basis in the brain.

To address this question, we analyzed blood oxygenation level dependent (BOLD) responses in the cortex of human subjects before and after they were trained on a motion direction discrimination task (Fig. 6A, ref. 37). Twenty-two human subjects participated in the motion VPL study. Subjects were trained for 10 days on a fine direction discrimination task, and psychophysical and fMRI tests were performed before and after training.

**Figure 6.**
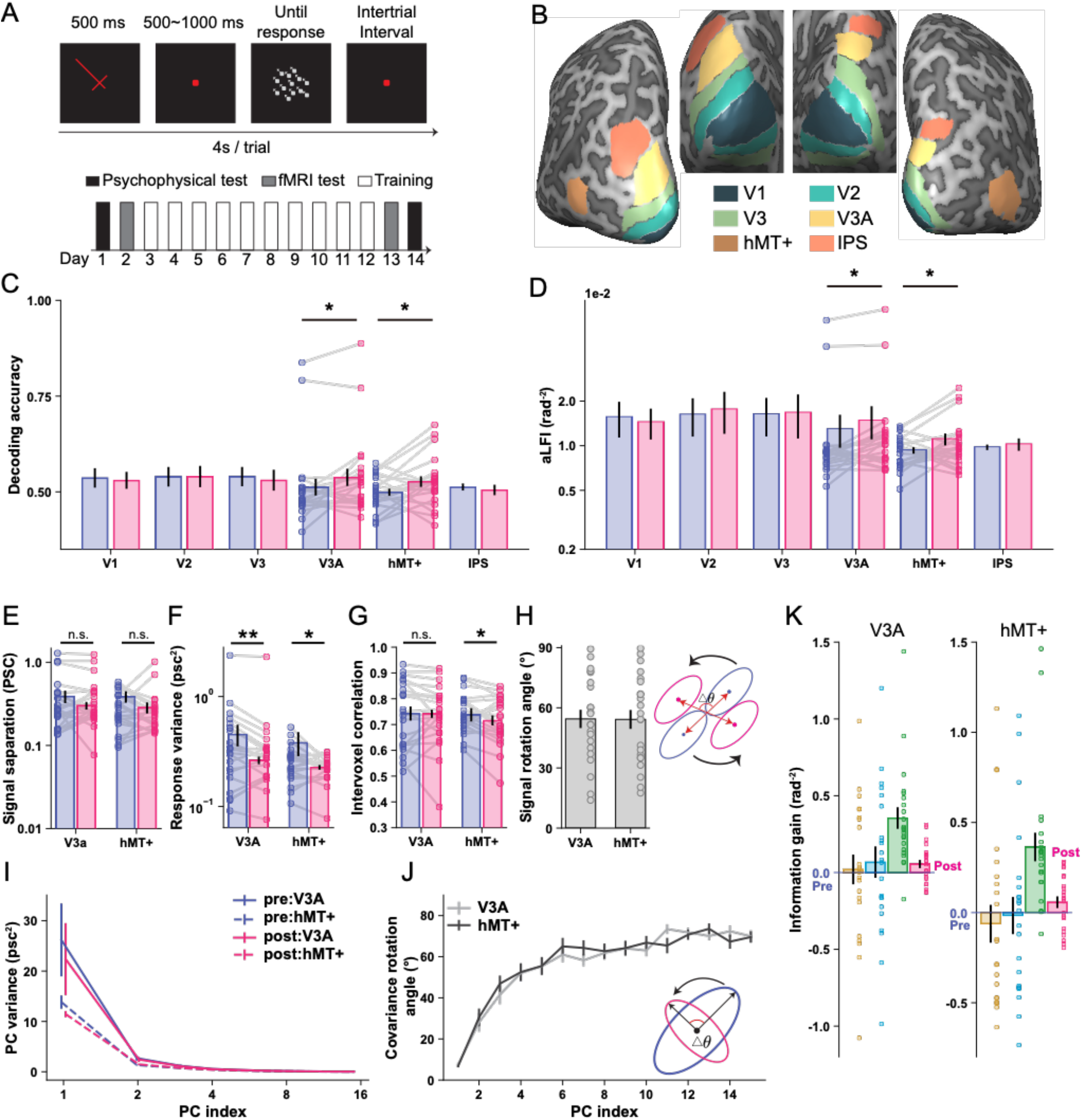
Trial diagram and training paradigm (***A***), and regions-of-interest in a typical subject (***B***). Motion direction discrimination training in humans significantly improves decoding accuracy (***C***) and averaged linear Fisher information (***D***) in areas V3A and hMT+, a finding consistent with several existing fMRI studies of motion VPL. Note that the four data points in V3A appear as outliers in ***C&D***, but the results still hold if these data points are removed. Motion direction discrimination training does not significantly change signal separation in V3A and hMT+ (***E***), but reduces voxel response variance in V3A and hMT+ (***F***) and intervoxel noise correlations in hMT+ (***G***). Similar to the motion DCNNs, Motion direction discrimination training in humans also rotates stimulus distributions (***H***), reduces the variance of high variance PCs (***I***), and warps the covariance directions (***J***). The patterns of information gain associated with the four mechanisms (***K***) are consistent with those in the DCNNs of both orientation and motion VPL. The unit “PSC” represents percent signal change of blood oxygenation level dependent signals. Individual data points represent the human subjects. Error bars in all panels represent the S.E.M. across subjects. Significance conventions are: *, *p* < 0.05; **, *p* < 0.01.

We identified the early visual areas (V1-V3), the motion-selective regions (V3A and hMT+), and the decision region (intraparietal sulcus, IPS) using independent functional localizer experiments (Fig. 6B). We estimated single-trial responses of voxels in these regions and then performed decoding analyses in these predefined regions, and found that motion training significantly enhanced decoding accuracy (Fig. 6C) and averaged linear Fisher information (Fig. 6D) in areas V3A and hMT+, a result consistent with several human fMRI studies on motion VPL 3,4,22.

We further investigated the coding principles in areas V3A and hMT+, and repeated the above analyses of DCNNs on fMRI data. Note that here we performed the same analyses on voxels instead of artificial neurons in DCNNs. Consistent with the predictions of the DCNNs, motion direction discrimination training in humans did not increase signal separation (Fig. 6E) but markedly reduced voxel response variance (Fig. 6F) in both areas. Motion direction discrimination training also significantly reduced intervoxel correlations in hMT+ (Fig. 6G). The mechanism of signal rotation was also evident, as indicated by the average ∼55º rotation of the signal vectors in both areas (Fig. 6H). In addition, training warped the magnitude and direction of the covariance (Fig. 6I&J). Most importantly, the respective contributions of these four mechanisms in both brain regions were similar to the pattern in the DCNNs (Fig. 6K).

### Neural geometric changes of contrast discrimination learning in monkey V4

So far, we have verified our neural geometry theory of in DCNNs and in the human brain. However, voxel responses in fMRI studies reflect macroscopic brain activity that aggregates the responses of ∼300, 000 – 50, 000 neurons 38. It remains unclear whether the mechanisms we have discovered so far also exist at the local circuit level of single cells or small clusters of cells. Although in the DCNNs our neural geometry theory generates two novel predictions that (1) visual training induces novel mechanisms (e.g., signal rotation and manifold warping) that can only be revealed by analyzing multivariate responses, and that (2) manifold shrinkage is the primary mechanism of VPL. To our knowledge, these predictions have not been systematically tested using intracranial recording.

To further test our hypotheses on neuronal spiking activity, we analyzed the population responses of V4 neurons in two monkeys (Fig. 7A) at the early and the stage of training in which they learned to perform on a fine contrast discrimination task (Fig. 7B, ref. 14). In this task, each monkey was presented sequentially with two identical Gabor patches with different contrast levels. The contrast of the reference (i.e., the first) stimulus was always fixed at 30%, and the contrast of the target (i.e., the second) stimuli varied systematically near the reference contrast (i.e., 27/28/29/31/32/33%). This contrast discrimination training significantly improved behavioral performance (Fig. 7C). Most importantly, responses of multiple channels were continuously recorded via chronically implanted electrodes in area V4 (21 and 29 channels for monkey 1 and 2, respectively) throughout training (21 and 23 training sessions for the two monkeys, respectively). This continuous multi-unit recording is the key to disentangling population-level changes associated with VPL.

**Figure 7.**
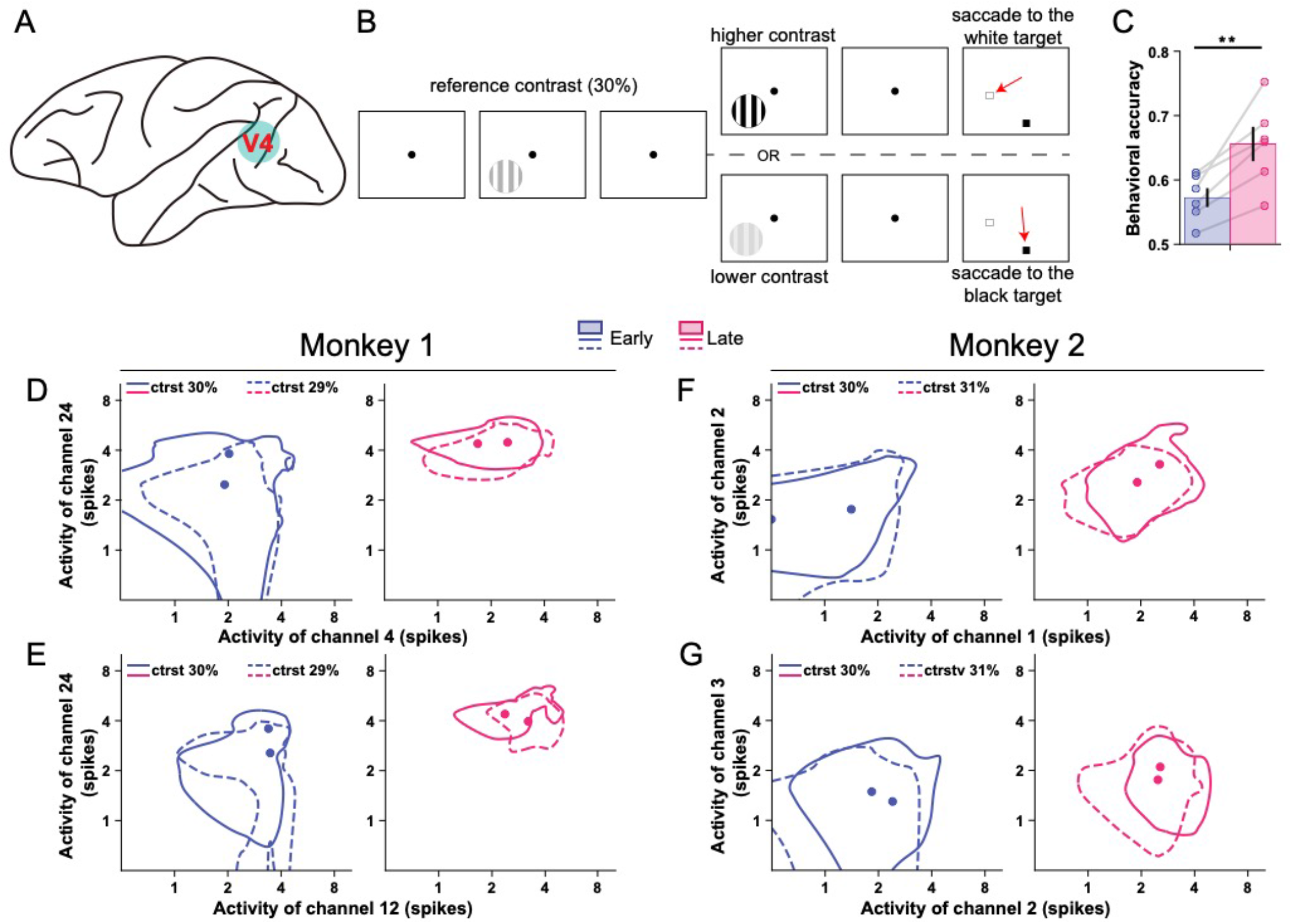
We analyzed population responses in area V4 (***A***) of two monkeys while they were trained on a fine contrast discrimination task (***B***). The first four and last four training sessions were grouped as pre- and the post-test conditions, respectively. Contrast discrimination training significantly improved behavioral performance from the early to late stage of training (***C***). All individual data points in ***C*** represent the six target contrast conditions (27, 28, 29, 31, 32, and 33%; the reference contrast is 30%). Each point is averaged over the two monkeys. See plots for individual monkeys in Supplementary Fig. S5. Error bars indicate the S.E.M. across the six conditions. ***D-G*** illustrates the full width at half maximum of the response distributions of four pairs of channels at pre- and post-test (***D&E*** for monkey 1 and ***F&G*** for monkey 2). Solid lines represent 30% reference contrast, and dashed lines represent 29% and 31% target contrast in monkey 1 and monkey 2, respectively. These results show that learning systematically changes the geometries of the multivariate responses.

We used the above analyses (previously applied to DCNNs and human fMRI data) and applied them to the monkey V4 responses, and again found highly consistent results. First, contrast discrimination training improved stimulus information at the population level (Fig. 8A&B). Second, at the individual level, contrast discrimination training also significantly reduced Fano factors (Fig. 8C) and noise correlations (Fig. 8D), consistent with several existing findings. Interestingly, while the trial-by-trial variance was significantly reduced after training (Fig. 8F), no apparent change in signal separation was observed (Fig. 8E), suggesting the predominant role of manifold shrinkage. Importantly, we again observed evidence for signal rotation (Fig. 8G) and manifold warping (Fig. 8H&I). The stepwise information analyses also qualitatively replicated the relative contributions of the four mechanisms to the total stimulus information encoded in the population (Fig. 8J).

**Figure 8.**
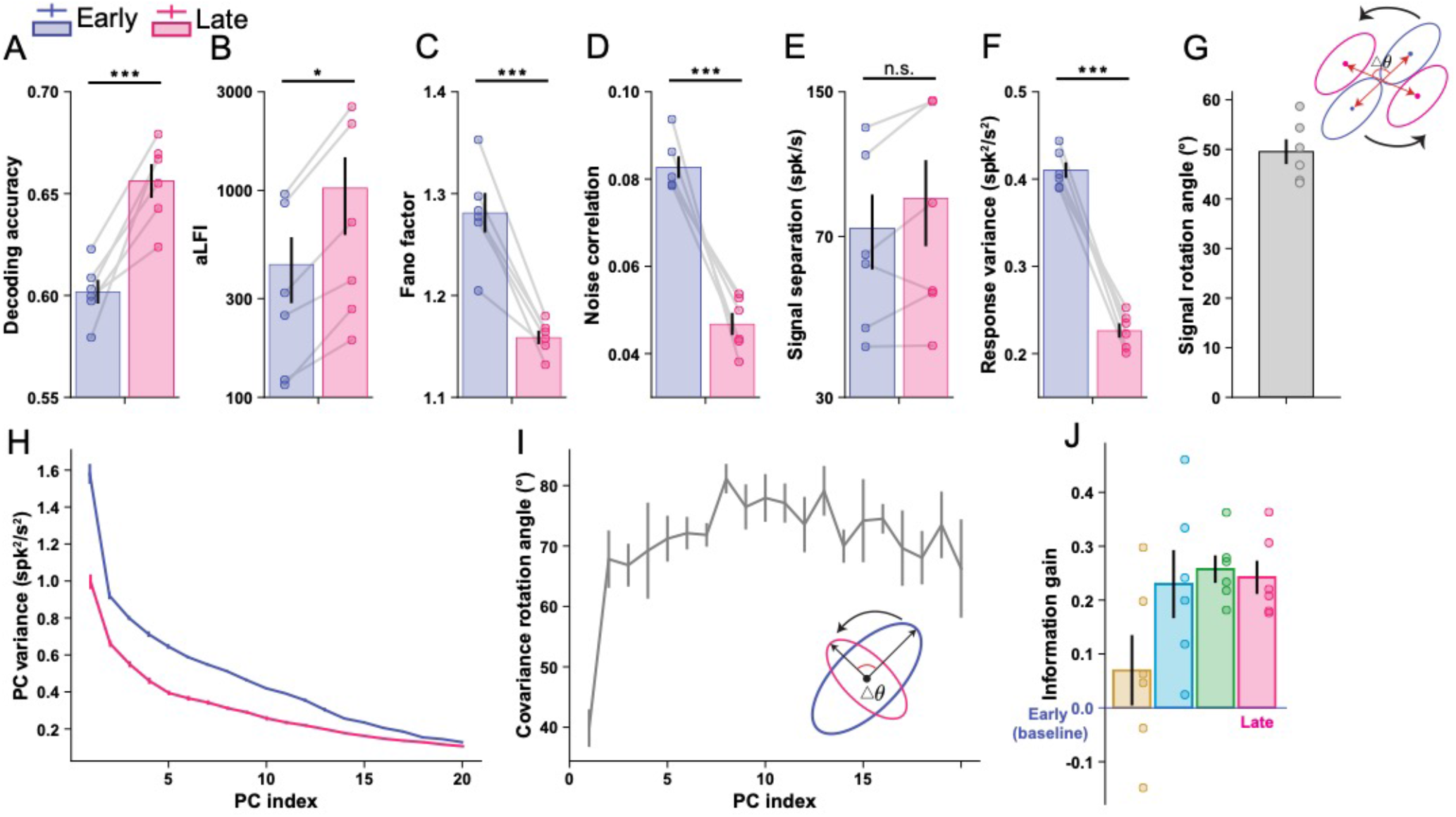
Population activity analyses of contrast discrimination learning in monkey V4. Contrast discrimination training significantly enhanced stimulus information at the population level (***A&B***). Consistent with VPL in the DCNNs and the human brain, training monkeys on a contrast discrimination task reduced Fano factors (***C***), noise correlations (***D***), and response variance (***F***), but had no significant effect on signal separation (***E***). We also found evidence for signal rotation (***G***) and manifold warping (***H&I***). The stepwise information analyses also show the similar pattern of the four mechanisms (***J***). The unit “spk/s” indicates the number of spikes per second (i.e., firing rate). We calculate aLFI and information gain using stimulus contrast as decimal values (i.e., 0.29), so they have arbitrary units. The definitions of the individual data points and error bars are the same as in Fig. 7.

Taken together, we proposed a neural geometry theory of VPL that links improved population representations to four possible neurocomputational mechanisms. We quantitively verified our theory across species, behavioral tasks, experimental conditions, measurement modalities, and brain regions. These converging results in humans, monkeys, and DCNNs suggest that our neural geometry theory can account for VPL in both biological and artificial systems.

## DISCUSSION

Our neural geometry theory bridges response properties at the single-unit level and those at the population level through geometric interpretations in a high-dimensional neural space. This theory allowed us to quantify possible contributions of single-unit responses to population representations. It has been controversial whether single-unit properties such as sharpened tuning curves 9,10 or reduction of noise correlations 11,12 contribute to VPL. Our information-theoretic analysis on neural geometry suggested that, although these properties were indeed observed, they did not contribute significantly to the improved population representations associated with VPL. Rather, we found that the totally overlooked mechanism—the response variance of individual units (i.e., manifold shrinkage)—is the primary contributor to the improved population representations associated with VPL. These results were further tested on deep convolutional neural networks, human fMRI data, and monkey neurophysiological data associated with different VPL tasks and brain regions.

Given the pronounced changes in tuning curves and changes in noise correlations observed after training, why do they not contribute to VPL? Conventional studies treat changes in tuning curves and in noise correlations as two independent factors mediating VPL. However, in our theory, the effects of tuning curve changes partially overlap with those of noise correlations (Eq. 4 in Methods and Materials). According to the neural geometry theory, the effects of tuning curve changes can be decomposed into two parts - signal enhancement independent of noise correlations and signal rotation interacting with noise correlations. We observed minimal evidence for signal enhancement (i.e., comparable magnitude of signal separation before and after training), and it certainly makes minimal contributions to population representations. Although we observed clear evidence for signal rotation and manifold warping, which correspond to the interactions between tuning curve and noise correlation changes, what matters here is the relative geometric relationship between the direction of noise correlations and signal vectors, rather than their absolute changes. Therefore, their respective contributions appeared significant but their overall joint effects were minimal because their effects canceled each other out (Supplementary Figure S6). Our findings challenge previous studies that have only assessed the effects of individual factors without considering the joint effects of multiple factors 11,13. The joint effects of multiple factors may be quite different from simply summing the effects of individual factors.

The novel finding that manifold shrinkage is the primary contributor to improved population representations is of unique significance in constraining the model of VPL. In manifold shrinkage, the total variance of the high-dimensional distributions is scaled down (i.e., *λ* in Eq. 4 is reduced), but the overall shape of the distributions remains unchanged. Thus, the two stimulus distributions simply shrink to a “smaller size” (Fig. 4). Note that manifold shrinkage is independent of any tuning changes and noise correlation changes. We also emphasize that manifold shrinkage and manifold warping are two different mechanisms. In our theory, manifold warping redistributes the variance of the high-dimensional distributions in different directions (i.e.,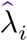 and *ξ*_*i*_ in Eq. 4 are changed) but, unlike manifold shrinkage, the total amount of variance remains unchanged. Thus, the shape of the two stimulus distributions is significantly warped.

Our neural geometry theory bridges existing human imaging and monkey neurophysiological studies by linking changes at both the individual and population levels. Most existing human imaging studies on VPL have focused only on improved representations at the population level 3,4,22, whereas most single-unit studies in monkeys can only measure changes in individual neurons 9,10. Only two empirical studies have attempted to address the relationship between the two levels using human fMRI 22 and monkey population recording 13. In these studies, researchers analyzed population responses by projecting high-dimensional neural manifolds onto a one-dimensional optimal decision plane. However, we argue that high-dimensional population responses, once projected onto a one-dimensional decision plane, no longer contain useful information about changes in individual units. In their approach, all changes in individual units are conflated (see our analytical derivations in Supplementary Note 1) and no longer distinguishable. In contrast, we evaluated all mechanisms within the high-dimensional neural manifolds *per se* without any dimension reduction operation. Our approach can explicitly disentangle and quantify the effects of individual factors (see similar analysis of our data in Supplementary Fig. S4). Our highly consistent findings in both artificial and biological visual systems demonstrate the effective utility of neural geometry theory in various domains of cognitive neuroscience. Although we only examined VPL of orientation, motion direction, and contrast discrimination, we argue that our neural geometry theory *per se* serves as a general framework to reveal whether the same as or different mechanisms than VPL in this study play a role in VPL for other tasks, possibly involving distinct brain regions. Neural geometry theory is also highly relevant for understanding the mechanisms of many other cognitive processes. For example, attention 39 and arousal 40 have also been shown to be associated with several neural changes (e.g., noise correlations). Similarly, it remains unclear how these effects contribute to improved population representations. Elucidating various cognitive processes from the perspective of neural geometry is a promising direction for future studies.

In summary, the long-standing controversy over whether changed tuning curves or reduced noise correlation, which are both property changes at the single-unit level, plays a significant role in VPL has been addressed by our research within the framework of neural geometry theory. Our converging results from monkey multi-unit recordings, human fMRI recordings, and neural network applications associated with various VPL tasks indicate that neither changed tuning curves nor reduced noise correlation affect the population response associated with VPL. Instead, previously overlooked mechanisms such as manifold shrinkage were identified as significant drivers of population response changes linked to VPL. This approach sheds light on how properties at single-unit level are linked to those at the population level in various types of VPL, as well as in other forms of learning and memory.

## METHODS AND MATERIALS

### DCNN modeling of orientation VPL

#### Stimuli

The network was trained to discriminate whether a target stimulus was tilted 1º clockwise or counterclockwise relative to a reference stimulus. All reference stimuli in the orientation discrimination task were Gabor patterns (227 × 227 pixels, spatial frequency=40 pixels/cycle, standard deviation of the Gaussian spatial envelope=50 pixels). The stimuli were varied in contrast (0.1 to 1.0 in 0.1 increments) and image noise level (8 levels: 0.005, 1, 5, 10, 15, 30, 50, 75). Similar to existing psychophysical studies 41, the image noise level is defined as the fraction of pixels randomly selected and replaced by Gaussian noise with a standard deviation of 15 gray level units. To mimic intrinsic sensory noise, we also added Gaussian white noise (standard deviation=10) to each stimulus 19. To match the spatial frequency of noise and signal, the size of the replaced pixels was set to be 8 × 8. Four reference orientations (35, 55, 125, and 145º) were used and we trained four DCNNs for each of the four reference orientations.

#### Neural networks and training

A DCNN 20 was used to simulate the orientation VPL. We retained the first five convolutional layers of the pre-trained AlexNet and replaced its three fully connected layers with a single linear fully connected layer for perceptual choice. The network was configured in a Siamese fashion to perform the two-alternative forced-choice (2AFC) task: the same network was fed with both the target and the reference stimuli, producing two scalar outputs, *h*_*t*_ and *hr*, respectively. The network then made the final decision with a probability *p* (classification confidence) calculated by the sigmoid function:

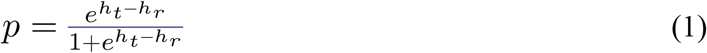

The entire training procedure consisted of two distinct phases: the pre-training phase and the VPL phase. In the pre-training phase, the network was trained on full-contrast noiseless stimulus pairs to understand the task and to establish the pre-test baseline. In the VPL phase, the network was trained on stimulus pairs across all contrasts (10 levels) and noise levels (8 levels). The network was trained for 5000 epochs in the pre-training phase and 500 epochs in the VPL phase using the stochastic gradient descent learning algorithm. The learning rate and the momentum were set to 1e^-5^ and 0.9, respectively. The parameters were updated to minimize the cross-entropy loss between the network outputs and the true stimulus labels. The initial parameters in the fully connected layer were set to zero, as in ref. 19, while those in the convolutional layers were taken directly from a pre-trained AlexNet available at http://dl.caffe.berkeleyvision.org/bvlc_AlexNet.caffemodel. We trained one model for each of the four reference orientations, and the entire procedure was repeated 10 times for each reference orientation to control for randomness. All model and training procedures were implemented using PyTorch 1.13.1.

#### Behavioral and neural changes

The behavioral performance of the network was evaluated by measuring its classification confidence (Eq. 1) at all combinations of contrast and noise before and after the training stage. Specifically, in each trial, the firing rate of each artificial neuron was measured as the output of local response normalization or ReLU layers, averaged over all locations. All measurements were obtained by simulating 1000 trials for better estimation. To ensure that units were truly driven by the stimuli, only units with a mean firing rate greater than 0.001 before and after training were included in the analyses 19.

The behavioral TvN curves of the model were derived for comparison with human psychophysical results. Specifically, for each noise level, the contrast threshold was obtained by interpolating the pre- and post-test accuracies of 55% and 70% respectively. To perform population decoding analyses, we trained a linear classifier on the firing rates of the artificial neurons to discriminate the target and the reference stimuli. The classifier was trained on half of the 1000 simulated trials, while the other half served as the test dataset.

To characterize the response properties of individual units, we measured orientation-selective tuning curves by sweeping the orientation of high-contrast stimuli from 0 to 180º. The tuning curves were derived by averaging 100 simulated trials for each orientation. The resulting tuning curves were then smoothed with a 10º Gaussian kernel. To control the heterogenous response range across units, we then normalized the tuning curves of each unit by its maximum response and averaged the tuning curves across units to obtain the group-level tuning curves. The group-level tuning curves were then fitted with a Gaussian function and rescaled to 0∼1 for better comparison.

To calculate the Fano factor of each unit, we simulated 1000 trials for each reference orientation. The Fano factor of each artificial neuron is defined as the ratio of the variance of the firing rate to its mean. Similarly, noise correlations between artificial neurons were calculated as the correlations between unit firing rates over the 1000 simulated trials for each reference orientation. We took the median of the Fano factor across units in each layer to generate the data plot (Fig. 2I). We took the median of the lower triangle of the noise correlation matrix in each layer to generate the data plot (Fig. 2J). The error bars in Fig. 2I&J represent the standard errors across four reference orientations.

#### Linear Fisher information analyses

To understand how neural activation contributes to behavioral improvements, we applied linear Fisher information analysis to population responses. We considered the firing rates of the same groups of units under the reference and the target stimulus conditions as two distributions in a high-dimensional neural space. We refer to the *signal vector* as the vector connecting the mean of the two distributions. A signal vector is calculated as the difference between the mean firing rates of units to two stimuli. The signal separation is referred to as the modulus length of the signal vector, and the angle of the signal vectors before and after training is referred to as the signal rotation angle.

To measure how much information was contained in a layer per unit, we calculated the average linear Fisher information (aLFI) as follows:

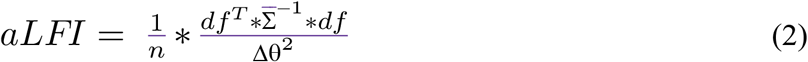

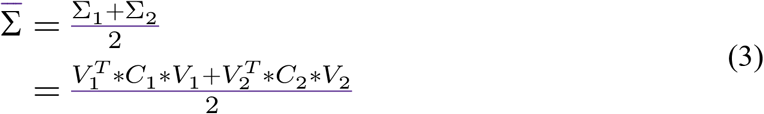

where *n* is the number of units in a layer, Δ θ is the separation between the target stimulus and the reference stimulus (i.e., 1º). *df* is the signal vector. 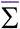 is the mean of the covariance matrices (i.e., Σ_1_ and Σ_2_) of units responding to the two stimuli. *V* is a diagonal matrix with the variance of the units as the diagonal terms. *C* is the correlation matrix of the population with all diagonal elements equal to 1.

To further elaborate on the potential mechanisms of the improved LFI, we performed an eigen decomposition on the covariance matrix 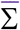, where we obtained *λ*_*i*_, the eigenvalue of 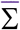 and *ξ*_*i*_, its corresponding normalized eigenvector. The aLFI can be rewritten as follows:

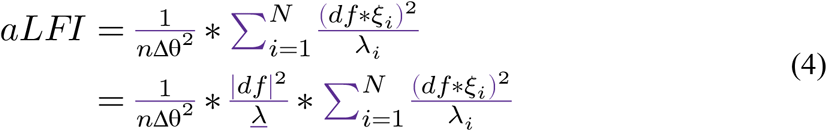

where *λ* is the mean variance, and 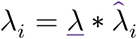. 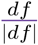is the unit vector with length of 1 and direction as the same as the signal vector *df*. According to Eq. 4, we disentangled the potential mechanisms of improved LFI into four subparts: signal enhancement, reflected by the modulus length |*df*|; manifold shrinkage, reflected by the mean variance of 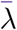; signal rotation, reflected by the direction of the signal vector *df*, and manifold warping, reflected by the relative angle of both *ξ*_*i*_ and 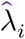. We applied a stepwise approach to assess their respective contributions by sequentially allowing only one mechanism to occur and calculating the resulting changes in *aLFI*. Specifically, we first calculated *aLFI* at pre-test as:

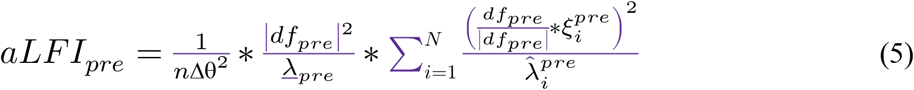

Only considering the effect of signal enhancement, we can calculate its effect as *aLFI*_*se*_:

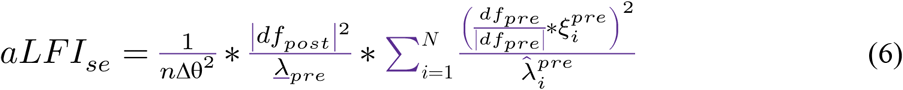

Note that the only difference here is that the 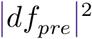 in Eq. 5 is replaced by the 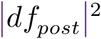 in Eq. 6. The difference between *aLFI*_*se*_ and *aLFI*_*pre*_ is considered as the information gain introduced by the signal enhancement mechanism (i.e., the brown bars in in Fig. 4B). Following this idea, we can calculate the stepwise *aLFI* by one-by-one considering the effects of manifold shrinkage *aLFI*_*ms*_, signal rotation *aLFI*_*sr*_, and manifold warping (*aLFI*_*mw*_ or *aLFI*_*post*_) as:

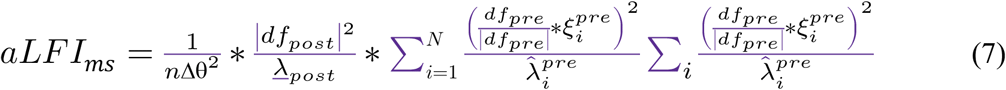

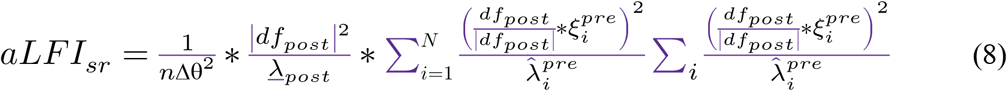

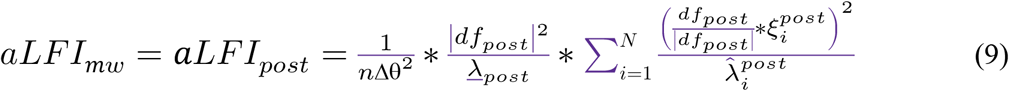

The information gain in Fig. 4B indicates the difference between *aLFI*_*se*_, *aLFI*_*ms*_, *aLFI*_*sr*_, and *aLFI*_*mw*_(i.e., *aLFI*_*post*_) as compared to the pre-test baseline . *aLFI*_*pre*_. They are shown as brown, blue, green, and magenta bars in Fig. 4B, respectively.

### DCNN modeling of motion VPL

#### Stimuli

The experiment used random dot motion (RDM) stimuli, which consist of a cloud of independent moving dots with some degree of coherence in a given moving direction 42. The network was trained to discriminate whether the moving direction of a target RDM stimulus was 4º clockwise or counterclockwise relative to its corresponding reference RDM stimulus. To meet the network’s specifications, the motion stimuli were 16-frame videos (112 × 112 pixels per frame). Within each frame, ∼100 dots were displayed, with each dot represented by a cross of 3 pixels in both height and width. We set 8 coherence levels (8.84, 12.5, 17.7, 25, 35.3, 50, 70.7, and 100%) and four reference directions (45, 135, 225, and 315º). The motion speed was 7.5 pixels/frame. All non-coherently moving dots appeared randomly in the image. The display of each frame was limited to a centered circle with a diameter of 112 pixels, with the surrounding areas displayed in black.

#### Neural network architecture and training

Our DCNN is a 3D convolutional neural network inherited from the C3D network for action recognition 36. The original C3D consists of 10 convolutional layers and 3 fully connected layers. The main difference between C3D and AlexNet is that C3D uses 3D convolutional kernels to process spatiotemporal information. We kept the first 6 convolutional layers from the pre-trained C3D and replaced the 3 fully connected layers with a fully connected layer that outputs a single scalar. The number of layers was chosen to (1) keep roughly similar number of parameters to the orientation DCNN, and (2) to roughly match the number of ROIs in the human neuroimaging experiment. Similar to the orientation DCNN, the motion DCNN was also configured in a Siamese fashion to perform the 2AFC task based on the sigmoid function.

Similar to the orientation DCNN, the entire training procedure consisted of two phases: the pre-training phase and the VPL phase. During the pre-training phase, the network was trained on full-coherence noiseless RDM pairs, while during the VPL phase, the network was trained on stimulus pairs across all coherence levels (8 levels). The network was trained for 1000 epochs in the pre-training phase and 2000 epochs in the training phase using stochastic gradient descent (SGD) with a learning rate of 1e^-7^, the momentum of 0.9, and a weight decay of 0.0005. The parameters were updated to minimize the cross-entropy loss between the network outputs and the true stimulus labels. The initial parameters in the fully connected layer were normally randomized while those in the convolutional layers were taken directly from a pre-trained C3D available at https://download.openmmlab.com/mmaction/recognition/c3d/c3d_sports1m_16x1x1_45e_ucf101_rgb/c3d_sports1m_16x1x1_45e_ucf101_rgb_20201021-26655025.pth. The entire procedure was repeated 10 times for each reference direction to control for randomness.

#### Behavioral and neural analyses

The behavioral performance of the network was also evaluated by its classification confidence (Eq. 1) at all coherence levels before and after the visual training phase. In addition, the firing rates of artificial neurons were measured on each trial as the output of the ReLU layers, averaged over all locations and time points. All measurements were taken over 1000 simulated trials. To ensure that units were truly driven by the stimuli, only units with a mean firing rate greater than 0.001 before and after training were included in subsequent analyses.

To perform decoding analyses, we trained a linear classifier on the firing rates of the artificial neurons to discriminate between the target and the reference stimuli. To assess the performance of the classifier, we split all trials half-half as training and test datasets, and used the average performance of the test-set. For comparison with the electrophysiological data, we calculated the Fano factor of each unit as the ratio of the variance of the firing rate to its mean, and the noise correlations as the correlation between the firing rates of units when viewing the same RDM stimulus. In addition, to measure how much information was contained in a layer per unit, we calculated the aLFI (see above).

We further validated the computational mechanisms in the motion direction discrimination task. To this end, the firing rates of the same group of units under the reference and the target stimuli were also considered as two distributions in a high-dimensional neural space. In the high-dimensional neural space, we defined signal vector, signal separation, variance, correlation, signal rotation angle, PC strength, and PC rotation angle as above.

Again, we computed linear Fisher information using a stepwise approach. For all models, we sequentially added signal enhancement, manifold shrinkage, signal rotation, and manifold warping to the calculation of linear Fisher information and see how the information within units varies with all four mechanisms. Fig. 5L shows the results of the stepwise analysis in layer 6. Supplementary Fig. S3 shows the results in all six layers of the motion DCNN.

### Human fMRI experiment

The human fMRI experiment data have been published in ref. 37 for different research questions. The core analyses in this study beyond preprocessing and ROI definitions are novel. We provide relevant methods as follows and more detailed methods in Supplementary Note 2 to avoid cross-referencing.

#### Subjects and Experimental procedures

A total of 22 human subjects (10 males and 12 females, ages 17-25) participated in the experiment. All participants had normal or correct-to-normal vision. None of the participants were aware of the study’s objectives. All participants provided written informed consent, and the study obtained approval from the local ethics committee at Peking University.

All subjects were trained on a direction discrimination task (Fig. 6A, see Supplementary Note 2 for apparatus and stimulus details). The whole experiment consisted of three phases: pre-test (2 days), training (10 days) and post-test (2 days). On day 1 at pre-test and day 2 at post-test, subjects were tested on direction discrimination around 45º and 135º (angular difference 4º, 120 trials for each direction) to assess their behavioral performance before and after training. Subjects were trained on the fine direction discrimination task for 10 days. Half of the subjects were trained at 45º and the other half at 135º (see training details in Supplementary Note 2). Training-induced behavioral improvements have been reported in our previous work 37.

To assess the neural changes induced by visual training, two identical fMRI sessions were performed on day 1 at pre-test and day 2 at post-test, respectively. In each fMRI session, subjects completed four runs of the motion direction discrimination task. Each run contained 30 trials for 45º and 135º (i.e., a total of 120 trials for each direction). Each run also contained 15 fixation trials and the trial order was randomized.

#### MRI data acquisition

All MRI data were acquired using a 12-channel phase array coil on a Siemens Trio 3T scanner at Peking University. The T1-weighted anatomical data with a resolution of 1 × 1 × 1 mm3 were collected for each subject. Echo-planar imaging (EPI) functional data were collected for the motion direction discrimination task, retinotopic mapping, and motion localizer experiments. EPI data were acquired using gradient echo-pulse sequences from 33 axial slices, covering the whole brain. The standard EPI sequence used for data acquisition was as follows: a repetition time of 2000 ms, an echo time of 30 ms, a flip angle of 90º, and a resolution of 3 × 3 × 3 mm3. The slice order was interleaved ascending.

In addition to the four runs of the motion direction discrimination task, we also collected one or two retinotopic mapping runs 52,55 and a motion localizer run 43 to define regions-of-interest.

#### MRI data analyses

In Brain Voyager QX, the anatomical data were transformed into the Talairach coordinate space. For all functional data, the first four volumes of each functional run were discarded to allow the longitudinal magnetization to reach a steady state. The functional data underwent several standard preprocessing procedures, including slice timing correction, head motion correction, spatial smoothing, temporal high-pass filtering (GLM with Fourier basis set at 2 cycles), and linear trend removal. Brain Voyager QX was also used to preprocess the data of the retinotopic mapping experiment and the motion localizer experiment. We used the standard phase-encoding method to define the retinotopic visual areas V1, V2, V3, and V3A 44,45. A generalized linear model (GLM) was then applied to the motion localizer data to define the motion-selective voxels (hMT+ and motion-selective voxels in IPS).

The functional data of the motion direction discrimination task were preprocessed using SPM12 (www.fil.ion.ucl.ac.uk/spm). The data were aligned to the first volume of the first run of the first session, corrected for acquisition delay, and then normalized to the MNI coordinate space using an EPI template. We used the GLMdenoise package developed in ref. 46 without evoking multirun denoise procedures to estimate the single trial activity of voxels.

#### Voxel population response analyses

We adapted the analysis previously used for artificial neurons in neural networks to the single-trial fMRI response estimates. To improve signal-to-noise ratio, we selected the 60 most responsive voxels in each ROI at pre-test. We first investigated which ROI was involved in motion VPL by measuring the discriminability between two different motion conditions (trained direction, e.g., 45º vs. untrained direction, e.g., 135º) before and after training. We trained a linear classifier on the fMRI data to discriminate between the two motion conditions. To assess the performance of the classifier, we performed a leave-one-trial-out cross-validation, and the average performance on the leave-out test trial was used as the discriminability measure. We also computed the average linear Fisher information (see equations above) between the 45º vs. 135º conditions to quantify stimulus discriminability. We found that motion direction discrimination training significantly improved stimulus discriminability in V3A and hMT+. Therefore, we only included V3A and hMT+ voxels in the subsequently analyses.

Similar to the analyses in the DCNNs, we defined the signal vector, the signal separation, the variance, the intervoxel correlations, the signal rotation angle, the PC strength, and the PC rotation angle in the multivoxel high-dimensional space using the same method defined above (Fig. 6). In addition, we applied the same stepwise analysis approach of calculating aLFI to the fMRI data (Fig. 6K).

### Monkey multi-unit recording experiment

Part of the monkey psychophysical and neurophysiological data have been published in refs. 14,47. These previous studies showed qualitatively similar results of the learning-induced reduction in Fisher information, Fano factor, and noise correlations via different analysis methods. Other results and analyses on the characteristics of population responses in this study (i.e., Figs. 7&8), especially the validation of signal rotation and manifold warping mechanisms, as well as the stepwise information analyses, are novel. We provide relevant methods as follows and more detailed methods in Supplementary Note 3 to avoid cross-referencing.

#### Ethics statement and Data collection

The Newcastle University Animal Welfare Ethical Review Board approved all procedures in this study. All experimental procedures were carried out in accordance with the European Communities Council Directive RL 2010/63/EC, the US National Institutes of Health Guidelines for the Care and Use of Animals for Experimental Procedures and the UK Animals Scientific Procedures Act. This study included two male monkey monkeys (5 and 14 years of age).

#### Experimental preparation

The surgical procedure is described in ref. 48 and Supplementary Note 3. The headpost and electrode implementations are also described in Supplementary Note 3. Briefly, in monkey 1, two 4 × 5 grids of microelectrodes were implanted in area V4; in monkey 2, one 5 × 5 grid was implanted in V4. These chronically implanted electrodes allowed us to record population activity in area V4 over the course of visual training. Importantly, we were able to record stably from a few small multi-unit clusters. The stability of the recording is shown in ref. 14. Stable recording of multi-channel neuronal activity allows analyses of changes in population responses induced by training.

#### Behavioral task and monkey training

The monkeys were trained in a contrast discrimination task in which subjects were asked to decide whether the contrast of a test stimulus was higher or lower as compared to that of a reference stimulus by making a saccade to one of two distinct locations (Fig. 7B). On each trial, the subject first kept fixation on the center-of-screen for 512ms. After 539 ms of fixation, a vertically oriented reference Gabor stimulus with 30% contrast was presented, centered at the V4 Receptive Field (RF) coordinates. The outer diameter of the Gabor stimulus was truncated at 16° for monkey 1 and 14° for monkey 2. Following the Gabor stimulus, monkey 2 experienced an interstimulus interval of 512 ms. In contrast, monkey 1 experienced a randomly chosen interstimulus interval, ranging from 512 to 1024 ms. During the interstimulus interval, only the fixation dot was presented. A test stimulus was then presented for 512 ms. This test stimulus was identical in size and orientation to the reference stimulus but differed in contrast, with the contrast level chosen pseudorandomly. The test stimulus was followed by another blank period of 512 ms during which only the fixation dot was visible. After the fixation dot, two target squares, one black and one white with a size of 0.5° in size, appeared to the left and right of the location where the reference and test stimuli were previously presented. The monkeys were cued to make a decision once the fixation dot disappeared. The monkeys were required to make a saccade to the white square within a 2° × 2° window if the test stimulus had a higher contrast than the reference stimulus. Conversely, they were expected to make a saccade to the black square if the test stimulus had a lower contrast than the reference stimuli. A correct saccade was rewarded with a fluid reward, while an incorrect saccade led to no reward and a 0.2 s timeout period.

The two monkeys were first trained on an easy version (target contrast 5% or 90%) of the contrast discrimination task. After they were fully familiar with the easy task, the target contrast increased from 2 to 8, 12, and 14 levels. The data correspond to the 14 levels of target contrast (10, 15, 20, 25, 27, 28, 29, 31, 32, 33, 35, 40, 50 or 60%; Supplementary Note 3). We only focus on target contrast levels (27, 28, 29, 31, 32, and 33%) near the reference contrast (i.e., 30%) according to the definition of linear Fisher information.

#### Dataset and preprocessing

We used chronically implanted Utah arrays to record spiking activity. We refer to small multi-unit neuronal clusters recorded from a given electrode as ‘channels’. Twenty-nine and Twenty channels were recorded in monkey 1 and monkey 2, respectively. These channels exhibited good responses (signal-to-noise ratio [SNR] >1) on over 80% of the recording sessions (see SNR computation in Supplementary Note 3). Baseline activity matching was performed between sessions for multi-unit activity (MUA) data in order to obtain comparable activity levels across sessions.

#### Behavioral and neural analyses

We noticed that the relationship between neural activity and discriminability can change drastically during the stimulus presentation period, and through training, the improvement in discriminability can also vary over the course of the training period. We chose the first 4 and the last 4 training sessions as the early and the late phase of training. This choice ensures an overall sufficient and comparable number of trials at both pre- and post-test for further analyses.

To determine the time window, we systematically varied the time window and trained a linear classifier to discriminate between the reference and target stimuli, and obtained its performance through 10-fold cross-validation. We chose the time window with the largest change in decoding accuracy between the reference stimulus (30% contrast) and the target stimuli (29% or 31% contrast). For monkey 1, the chosen time window was 30-130 ms after stimulus onset. For monkey 2, the time window was 130-230 ms after stimulus onset. Note that this choice aims to maximize training effects on population representations (similar to the decoding analyses for first identifying V3A and hMT+ as the ROIs where learning effects are most pronounced in the human fMRI study), but does not guarantee the underlying mechanisms such as signal separation enhancement and manifold shrinkage. Also, varying the time window did not qualitatively change our results. We used a simple multivariate Poisson log-normal (MPLN) model (Supplementary Note 3, see also refs. 49-52) to estimate the trial-by-trial variability of population firing rates. We further use the estimated firing rates and covariance to compute all neural metrics mentioned above. We report all results in Figs. 7&8 for visual comparison with the DCNN and fMRI results above.

## ACKNOWLEDGEMENT

We thank Shuguang Kuai, Duje Tadin, Oh-Sang Kwon for valuable comments on the manuscripts. This work was supported by the National Natural Science Foundation of China (32100901 to R.-Y.Z, 3230085 to K.J.) and Natural Science Foundation of Shanghai (21ZR1434700 to R.-Y.Z), The monkey work was supported by the Medical Research Council, UK, G0700976.

## AUTHOR CONTRIBUTIONS

Y-A. C. and R-Y.Z. conceived and designed the study. Y-A. C. implemented the neural networks. J.K., S.L., and F.F., prepared and provided the preprocessed fMRI data. A.T., X.C., and M.S. recorded, organized, and preprocessed the monkey physiological data. Y-A. C. and R-Y.Z. performed in-depth analyses on neural networks, human fMRI data, and monkey electrophysiological data. Y-A. C. and R-Y.Z. wrote the first draft of the manuscript. All authors revised the manuscript and provided valuable feedback to the final manuscript.

