## Supplemental Information for "A neural geometry theory comprehensively explains apparently conflicting models of visual perceptual learning"

<u>**Table of Content**</u>

**Supplementary Figure S1**: Neural consequences of orientation VPL in all five layers.

**Supplementary Figure S2**: The orientation DCNN replicates the neuronal response properties in the primate early visual cortex.

**Supplementary Figure S3**: Neural consequences of motion VPL in all six layers.

**Supplementary Figure S4**: Replications of macaque neurophysiological and human fMRI experiments on signal enhancement.

**Supplementary Figure S5**: Neural consequences of contrast discrimination learning in two monkeys.

**Supplementary Figure S6:** The effects of the mechanism order in the stepwise analyses.

**Supplementary Note 1**: Analytical relation between the high-dimensional population responses and one-dimensional decision unit responses.

**Supplementary Note 2**: Supplementary methods for the human fMRI experiment.

**Supplementary Note 3**: Supplementary methods for the macaque multielectrode recording experiment.

**Supplementary Figure S1**

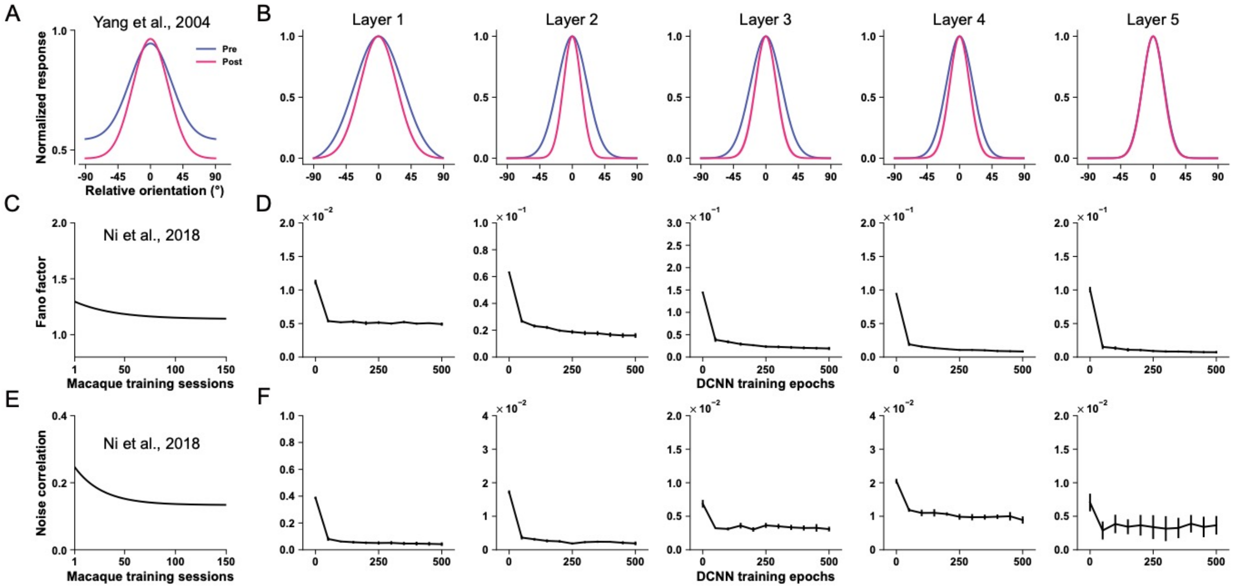


**Figure S1**. Tuning curve, Fano factor, and noise correlation changes in all five layers of the orientation DCNN. Perceptual training on the DCNN reduces the width of the tuning curves in layers 1-4. Perceptual training also reduces Fano factor and noise correlations in all five layers.

**Supplementary Figure S2**

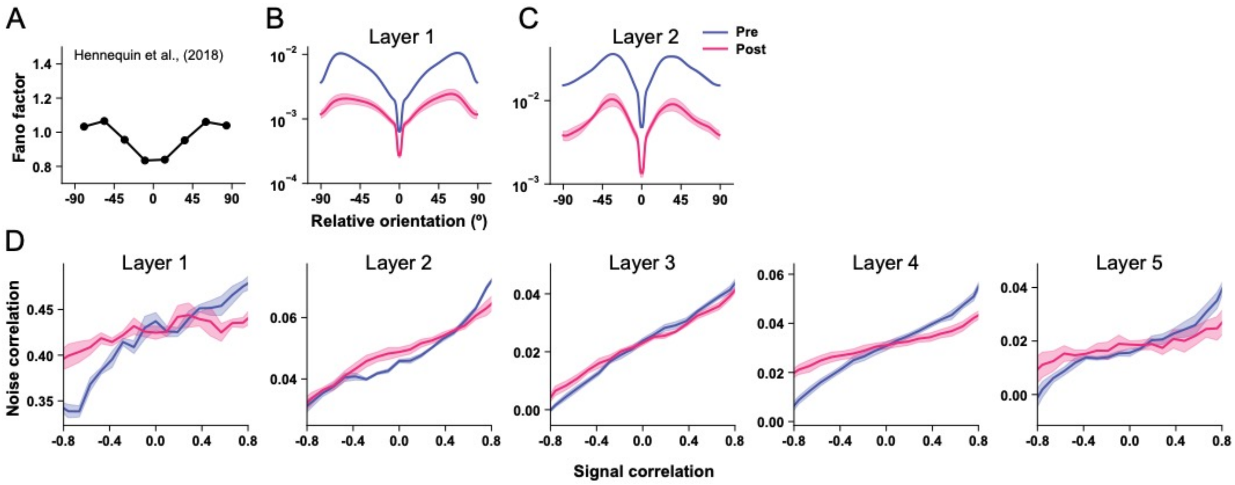


**Figure S2**. The DCNN of orientation VPL qualitatively replicates the relationship between orientation preferences and Fano factor in macaque V1 (***A***, reproduced from the datapoints from ref. 1). Here, the results of layers 1(***B***)&2(***C***) are plotted. The network also reproduces the positive relationship between signal correlations and noise correlations in the human 2,3 and macaque 4 early visual systems (***D***). Importantly, interneuron noise correlations are not uniformly reduced by training. In contrast, perceptual training reduces the noise correlations between neurons with similar tuning preferences (i.e., signal correlation > 0) and vice versa between neurons with opposite preferences (i.e., signal correlation < 0). These results are consistent with several theoretical and empirical findings on noise correlation changes induced by learning and attention 5-8.

### Supplementary Figure S3

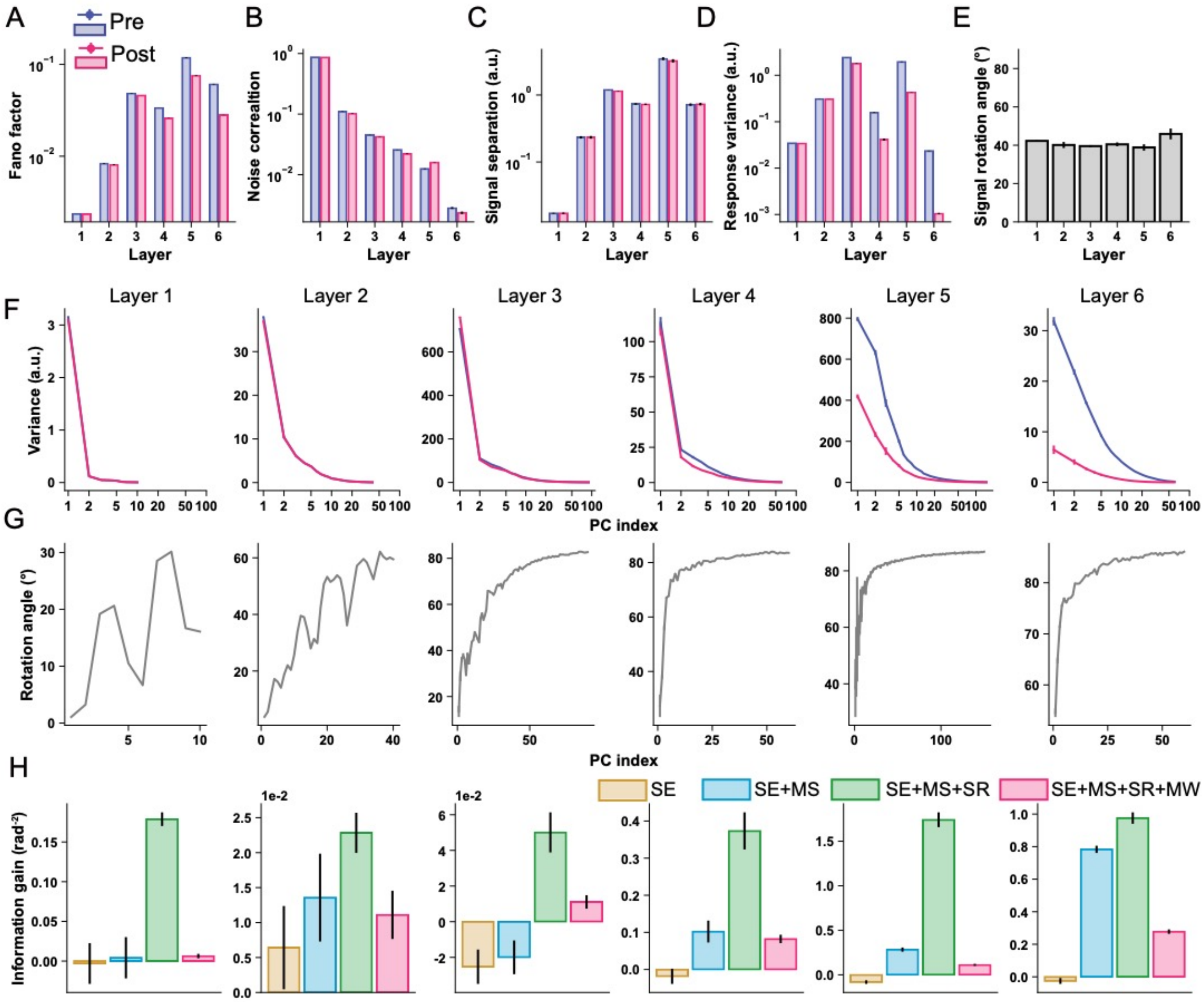


**Figure S3.** The results of all layers in the motion DCNN. The effects of motion direction discrimination training on Fano factor (***A***), noise correlations (***B***), signal separation (***C***), response variance (***D***), and signal rotation angle (***E***) in the DCNN model. The effects of training induced manifold warping in all six layers are shown in (***F&G***). The results of the stepwise information analyses are shown in (***H***). The figure conventions are the same as in the main text.

**Supplementary Figure S4**

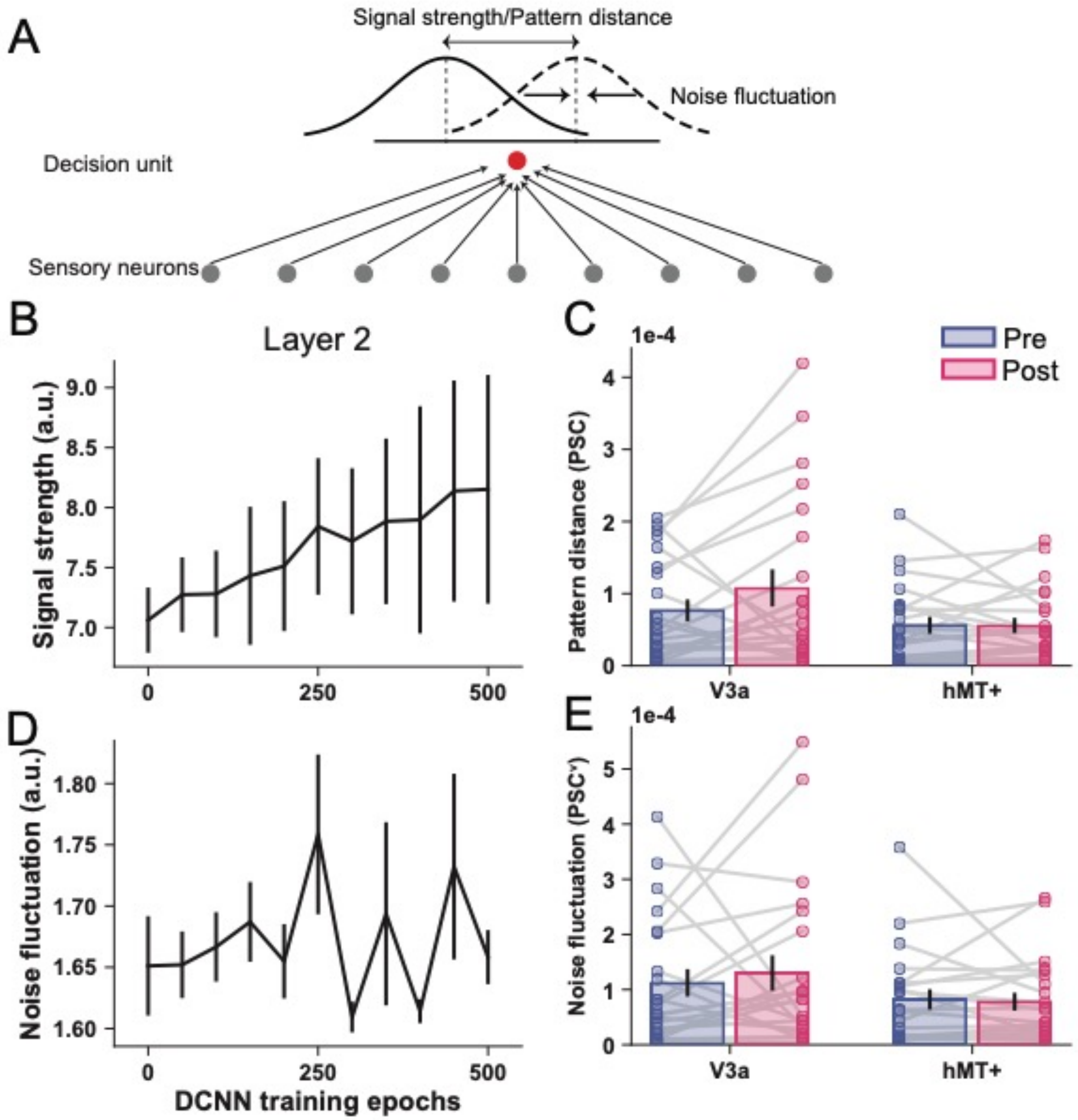


**Figure S4**. We applied similar analyses in refs. 9,10 to our data. Here, we project the multivariate population responses onto an optimal decision unit and derive the two 1-d activity distributions of the decision unit for the two stimuli. We replicate the results that, after this linear transformation, the effects of VPL appear to be signal enhancement in layer 1 of the orientation DCNN (***A***), and in V3A but not in hMT+ in the human brain (***B***). In contrast, noise fluctuation in layer 1 (***C***) and in V3A and hMT+ in the human brain (***D***) do not significantly increase. These results are consistent with the Fig. 6C in ref. 9 and Fig. 6C in ref. 10.

**Supplementary Figure S5**

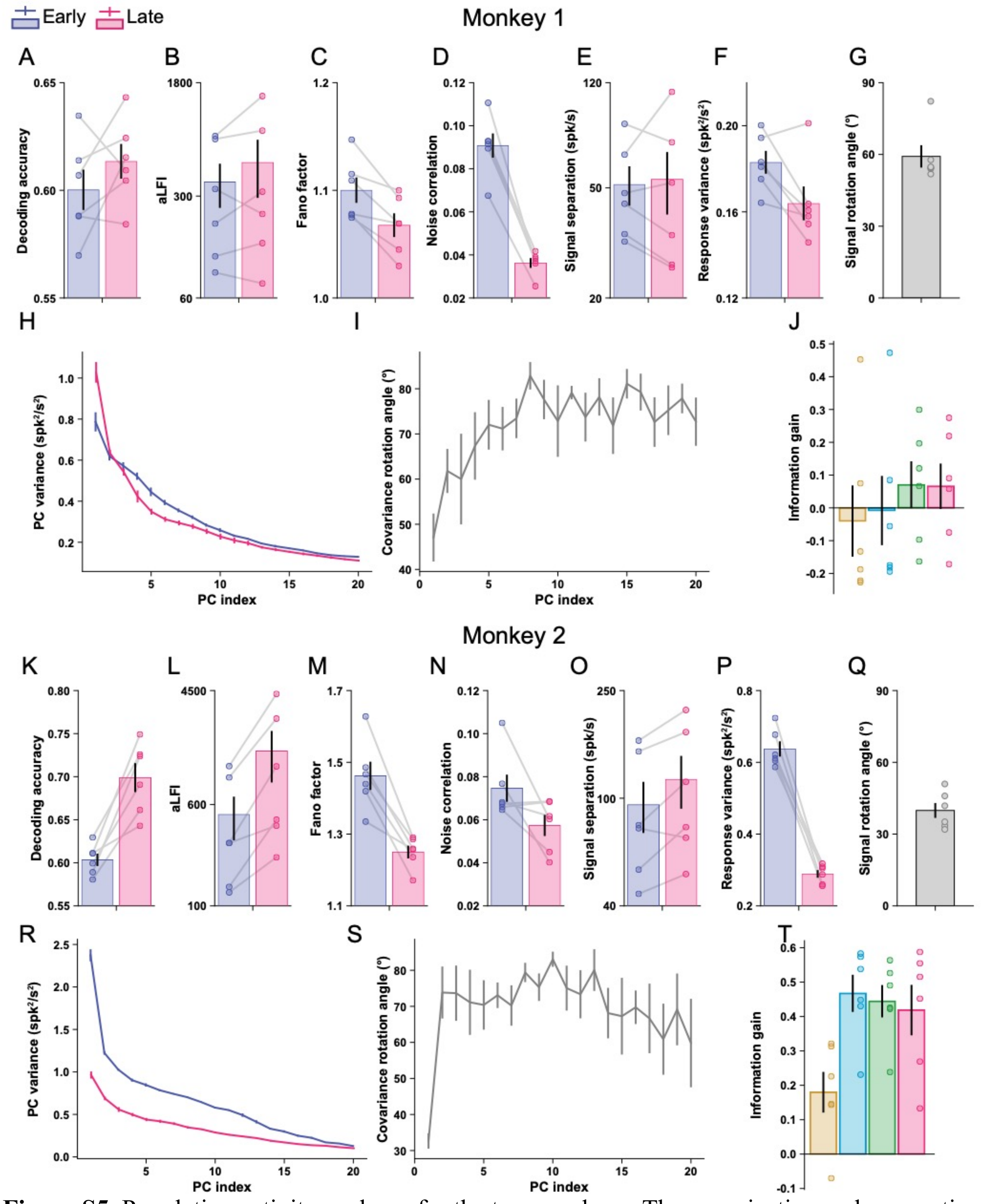


**Figure S5**. Population activity analyses for the two monkeys. The organization and conventions here are the same as Fig. 8 in the main text, except that we split the results for the two monkeys.

**Supplementary Figure S6**

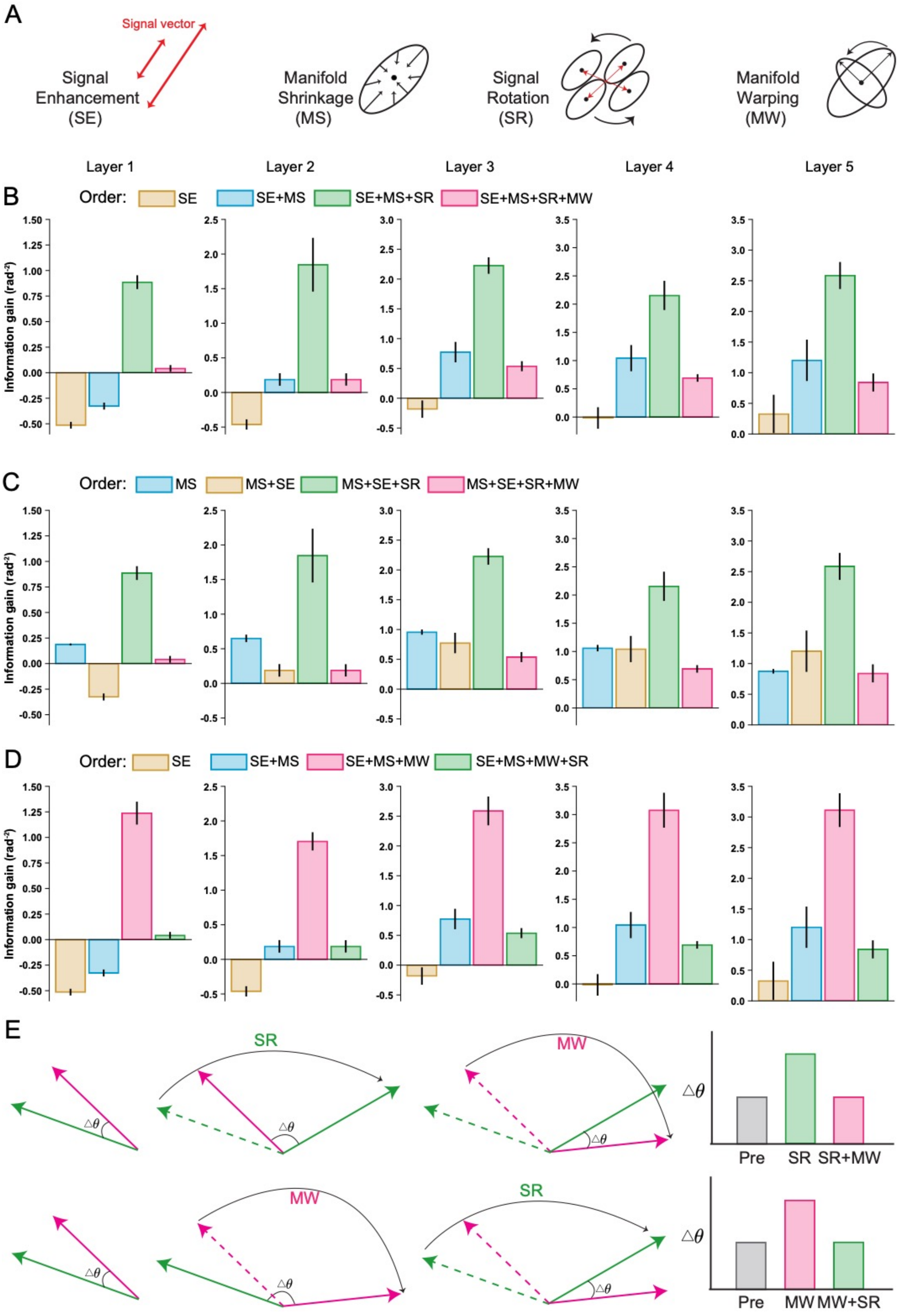
A
Signal vector
Signal Enhancement (SE)
Manifold Shrinkage (MS)
Signal Rotation (SR)
Manifold Warping (MW)
Layer 1
Layer 2
Layer 3
Layer 4
Layer 5
B
Order: SE SE+MS SE+MS+SR SE+MS+SR+MW
Information gain (rad⁻²)
C
Order: MS MS+SE MS+SE+SR MS+SE+SR+MW
D
Order: SE SE+MS SE+MS+MW SE+MS+MW+SR
E
SR
MW
Δθ
Pre SR SR+MW
Pre MW MW+SR

69

**Figure S6.** The effects of mechanism order on the results of stepwise information analyses. ***A&B***. The cartoon illustration of the four mechanisms (***A***) and the results of the stepwise information analysis according to the order of SE/SE+MS/SE+MS+SR/SE+MS+SR+MW, the same as in Fig. 4 in the main text. ***C***. The results of the stepwise information analysis according to the order of MS/MS+SE/MS+SE+SR/MS+SE+SR+MW. Here, the mechanisms of SE and MS are independent. Thus, changing their order does not change the positive effects of MS and the minimal or even negative effects of SE. More precisely, the information gains of SE and MS are quantitatively identical in ***B*** and ***C***. For example, in layer 1 of ***B***, the height of the yellow bar (~-0.5) indicates the negative effect of SE, which is equal to the decrement from the blue to the yellow bar in layer 1 of ***C***. Similarly, the increment from the yellow to the blue bar in layer 1 of ***B*** indicates the positive effect of MS, which is equal to the height of the blue bar in layer 1 of ***C***. ***D***. Compared to ***B***, we only switch the order of SR and MW in ***D*** (i.e., the results of SE/SE+MS/SE+MS+MW/SE+MS+MW+SR). The apparent effects of SR and MW appear to be opposite in ***B*** and ***D***. This is because what matters here is the projection of the covariance onto the signal vector, or in other words, the angle between the signal vector and the covariance direction [3,8,11,12]. Intuitively, the larger the angle, the more information the population contains. We explain this effect in cartoon illustrations in ***E***. No matter which mechanism (i.e., SR or MW) is considered first, it increases the angle between the signal vector and the covariance direction (i.e., increases information), but this effect is canceled out by further including the other mechanism. Thus, the apparently opposite effects of SR and MW actually reflect the same underlying mechanism.

**Supplementary Note 1: Signal enhancement and manifold shrinkage in multivariate sensory neural populations and one-dimensional decision unit**

In this section, we theoretically derive the relationship between the analyses of population representations and the 1D responses of the decision unit. Assuming $N$ sensory neurons in a population, we denote the mean firing rates towards the reference and the target stimuli as $\vec{\mu}_{ref}$, and $\vec{\mu}_{tgt}$, their covariance as $\Sigma_{ref}$ and $\Sigma_{tgt}$. We define the optimal linear decoder as $\vec{w}$. The decoder linearly transforms the high-dimensional population responses onto a decision plane to form the activity of the decision unit. The mean firing rates of the decision unit towards the two stimuli should be $\vec{w} \bullet \vec{\mu}_{ref}$ and $\vec{w} \bullet \vec{\mu}_{tgt}$ respectively while the variance should be $\vec{w}^T \Sigma_{ref}\ \vec{w}$ and $\vec{w}^T \Sigma_{tgt}\ \vec{w}$. According to the signal detection theory, we can define the separation between the two 1-d decision distributions to be the difference between the classes over the variance within the classes as follows:

$$J(\vec{w}) = \frac{(\vec{w}\bullet\vec{\mu}_{ref} - \vec{w}\bullet\vec{\mu}_{tgt})^2}{\vec{w}^T\Sigma_{ref}\ \vec{w} + \vec{w}^T\Sigma_{tgt}\ \vec{w}} = \frac{(\vec{w}\bullet(\vec{\mu}_{ref} - \vec{\mu}_{tgt}))^2}{\vec{w}^T(\Sigma_{ref}+\Sigma_{tgt})\ \vec{w}} \tag{S1}$$

In a linear discriminant analysis, the objective is to find a $\vec{w}$ that maximize $J(\vec{w})$, when $\vec{w} = k(\Sigma_{ref} + \Sigma_{tgt})^{-1}(\vec{\mu}_{ref} - \vec{\mu}_{tgt})$. The Eq. S1 can be further written as follows:

$$J(\vec{w}) = \frac{(\vec{w}\bullet(\vec{\mu}_{ref} - \vec{\mu}_{tgt}))^2}{\vec{w}^T(\Sigma_{ref}+\Sigma_{tgt})\vec{w}} = \frac{k\left(\vec{\mu}_{ref}-\vec{\mu}_{tgt}\right)^T(\Sigma_{ref}+\Sigma_{tgt})^{-1}\left(\vec{\mu}_{ref}-\vec{\mu}_{tgt}\right)}{\sqrt{k^2\left(\vec{\mu}_{ref}-\vec{\mu}_{tgt}\right)^T(\Sigma_{ref}+\Sigma_{tgt})^{-1}\left(\vec{\mu}_{ref}-\vec{\mu}_{tgt}\right)}} = \frac{LFI}{\sqrt{LFI}}, \tag{S2}$$

Note that after the linear transformation, the numerator—the difference of the means (signal separation defined in ref. 9)—is proportional to the linear Fisher information calculated from the sensory neuron population. The denominator—the square root of the variance of the two decision distributions (noise fluctuations defined in ref. 9)—is proportional to the squared root of the linear Fisher information. Therefore, their definitions of signal and noise actually reflect all four possible mechanisms identified in our main text.

More importantly, we also applied the similar analyses as in ref. 9 to the population responses in layer 1 of the orientation DCNN. We replicated the results reported in ref. 9. We also applied the same analyses to the voxel response data in the human fMRI experiment, and replicated the Figure 6 in ref. 10. In summary, we analytically derive the relationship between the signal/noise in high-dimensional population responses and the signal/noise in the one-dimensional responses of the decision unit. The two seemingly opposite conclusions can coexist in both artificial and biological visual systems.

**Supplementary Note 2: Supplementary methods for the human fMRI experiment**

_Experimental stimuli and procedures_. The random dot motion stimuli were presented to the participants on different display devices. In the behavioral sessions, a 40 cm wide CRT monitor with a resolution of 1024 × 768 pixels and a refresh rate of 60 Hz was used. In addition, a 48 cm wide LCD projector with a resolution of 1024 × 768 pixels and a refresh rate of 60 Hz was used to display the stimuli in the fMRI sessions. Stimuli were generated and displayed using Psychtoolbox 3.0 in the MATLAB environment. To ensure consistency, the viewing distance in the behavioral sessions was set at 75 cm from the screen. The method for generating the random dot stimuli followed the approach described in ref. 13. All stimuli were presented within an imperceptible aperture of 10º diameter centered on a black background. At any given moment, the aperture contained 400 dots, each moving in unison in the same direction at a speed of 4 º/s. Dots that moved out of the aperture were repositioned on the opposite side of the aperture to maintain a constant dot density.

At the beginning of each trial, a red reference cross was displayed for 500 ms. The orientation of the long arm of this red cross served as the reference direction for the subsequent motion stimulus. Following the reference cross, a red fixation dot appeared on the screen and remained there for the duration of the trial. After an unpredictable delay of 500 to 1000 ms, the motion stimulus was presented for 1500 ms or until participants responded. During the trial, participants were asked to indicate whether the motion stimulus rotated clockwise or counterclockwise relative to the orientation of the long arm of the reference cross. They were required to respond by pressing one of two designated keys according to their chosen direction preference.

The overall experimental design consisted of three distinct phases: the pre-test phase (2 days), the training phase (10 days), and the post-test phase (2 days). Prior to the pre-test phase, participants underwent a practice session consisting of 80 trials with an angular difference of 8º to familiarize themselves with the task. Following this practice session, baseline performance was measured for each participant using a 4º angular difference. This baseline assessment was performed using four blocks of 30 trials each, with reference directions set at 45º and 135º (90º representing the vertical upward direction). The pre-test functional magnetic resonance imaging (fMRI) session was conducted on day 2. Each participant completed four runs of the direction discrimination task, with each run consisting of 60 task trials (30 trials for each direction of 45º and 135º) interspersed with 15 fixation trials. The order of these trials was pseudorandomized within each run and across participants, except that the first two trials and the last three trials in each run were designated as fixation trials. In addition, functional localizer runs were conducted within the same session (see ROI definitions).

The behavioral training phase lasted 10 days and was designed to ensure that participants reached ceiling performance. On each training day, participants completed 10 training runs, each run consisting of 60 task trials and 10 fixation trials. The total duration of each training session was approximately 1 hour. Training began with an initial angular difference of 4 º, and this angular difference remained fixed for each training session. After reaching a minimum accuracy of 79.4% on the preceding day, the angular difference for the following day was reduced to one of the predetermined options (3, 2, 1.5, or 1º) in order to increase task difficulty. This procedure was designed to keep participants engaged and challenge them with increasingly difficult trials. A fixed set of difficulty levels was used instead of staircase procedures to ensure a consistent level of task difficulty. This approach resulted in the creation of more challenging trials over time. Half of the participants were trained on 45º stimuli, while the other half on 135º stimuli. During training, auditory feedback was provided when participants made incorrect responses. In addition, visual feedback in the form of "slow down" or "hurry up" prompts was presented when the response time was faster than 250 ms or slower than 1500 ms, respectively. These feedback mechanisms were implemented to help participants refine their performance during the training process.

The procedure of the post-test phase was identical to the pre-test phase, except that the fMRI session was conducted before the behavioral session.

*MRI data acquisition*. The fMRI sessions were conducted on Day 2 at the pre-test phase and on Day 1 at the post-test phase. On Day 2 at pre-test, besides the four task runs, we also collected 1 run of T1 images, 1 run of motion localizer, 1-2 retinotopic runs. The retinotopic runs used the standard phase encoding method 14. The motion-responsive voxels (V3A and hMT+) were identified with a localizer procedure 15. We presented random dot stereograms, which were static for 24 s and then traveled toward and away from the fixation for 8 s. The moving/stationary cycle repeated nine times. The size of the stimulus aperture was the same as that used in the main experiment.

**Supplementary Note 3: Supplementary methods for the macaque multielectrode recording experiment**

Because we analyzed preprocessed monkey data, most of the experimental procedures have been published previously in detail. We provide more method details here only to avoid cross-referencing. To preserve as much of the method details as possible, we duplicate most of the noncritical text from the original paper 7 for reference purposes only. All duplicated text is italicized.

*Surgical, headpost implementation, and electrode implementation procedures*. We duplicated most of the text as italics from the original paper:

*Details of surgical procedures and postoperative care have been published elsewhere* 16. *An initial surgical operation was performed under sterile conditions, in which a custom-made head post (PEEK, Tecapeek) was embedded into a dental acrylic head stage.*

*For surgical preparation, animals were sedated with ketamine. During surgery, anesthesia and analgesia were maintained by sevoflurane (gaseous, 1–3%) and alfentanil (intravenous 156 µg/kg/h), respectively. Blood pressure, rectal temperature, blood oxygen saturation and end tidal* $CO_2$ *were measured continuously. After the surgery, analgesic (metacam 0.1/kg) and prophylactic antibiotics (cephorex 0.5 ml/kg) was given for 3–5 days.*

*During surgery, the animals were placed in a stereotaxic head holder and the skull overlying the occipital and posterior temporal cortices was exposed. A craniotomy was made to remove the bone overlying V1, V2 and dorsal V4, using a pneumatic drill. The bone was kept in sterile 0.9% NaCl for refitting at the end of the surgery. The dura was opened to allow access to V4. Microelectrode chronic Utah arrays, attached to a CerePortTM base (Blackrock® Microsystems, connection dimensions of 16.5 mm [height] × 19 mm [base diameter] × 11 mm [body diameter]), were implanted under sterile conditions in the cortex, using a Blackrock microarray inserter. In monkey 1, two 4 × 5 grids of microelectrodes were implanted in area V4; in monkey 2, a 5 × 5 grid was implanted in V4. Electrodes were 1 mm in length, and their tips reached depths of up to 1 mm. Wire bundles were held in place with biologically compatible glue (histoacrylic), and the connector (CerePortTM) was secured to the skull with titanium bone screws. Following array insertion, the Dura was re-sutured over the array, the exposed area was thinly covered with sterile Tisseel Lyo two-component fibrin sealant (Baxter Healthcare), and the bone flap was reinserted into the skull (before the Tisseel had fully set). The bone flap was cross bridged to the surrounding skull using Synthes orbital plate fragments and Synthes titanium bone screws.*

*The electrode arrays were inserted under visual control into the gyrus between the lunate sulcus and the superior temporal sulcus. The recording locations were confirmed to be in area V4 in both animals via visual inspection immediately postmortem and by analysis of postmortem Nissl-stained brain sections.*

<u>Behavioral task and training procedures</u>. More detailed descriptions of task and training procedures are duplicated as italics from the original paper.

*Each monkey was trained in a contrast discrimination task in which he differentiated between two successively presented stationary Gabor gratings based on their relative contrasts (Fig. 7B).*

*Monkeys were initially trained on a very basic version of the contrast discrimination task at a location in the upper visual field, i.e., at a substantial distance from the RFs covered by our electrodes, which were located in the lower left visual field (for details, see below). When the animal understood the main concept of the task in the upper visual field, the stimuli were shifted to the left lower visual field. The stimuli (Gabor gratings, $\sigma = 4°$, spatial frequency = 2 cycle per degree, orientation = 90° vertical) were initially presented at an azimuth of −5° and an elevation of −16° in both monkeys (left and bottom compared to the fixation point). These coordinates covered the V4 RFs.*

*Each trial was initiated when the monkey held a touch bar and fixated on a small fixation spot (diameter = 0.1°, fixation window = 2° × 2°) which was presented on a grey background (52.17 cd/m$^2$). During the trial, if the monkey broke fixation before saccade cue onset or failed to respond within 1000 ms of the onset of the saccade cue, the trial was terminated immediately and followed by a 0.2 s timeout. We used different inter-stimulus intervals in the two animals for the following reason. We started training and recording in monkey 1, before doing so in monkey 2. We initially reasoned that a variable test onset would increase the animal's focus and thereby possibly learning. In monkey 2, the variable onset during the very basic training resulted in too many early trial abortions, which quickly vanished when we used a fixed delay. We therefore decided to use a fixed delay in that animal.*

*After monkeys performed well in this easy version of the task, the number of test contrasts was increased to 8 (5, 10, 20, 25, 35, 40, 60 and 90% contrast, on day 1 of the proper contrast discrimination task), then to 12 (10, 15, 20, 25, 27, 29, 31, 33, 35, 40, 50 and 60% contrast, on day 2 of the proper contrast discrimination task) and to 14 (10, 15, 20, 25, 27, 28, 29, 31, 32, 33, 35, 40, 50 or 60% contrast, from day 3 of the proper contrast discrimination task). In order to motivate subjects to complete each trial and discourage them from guessing on difficult trials, stimulus drumming was carried out using the 'repetition with delay' function on CORTEX following error trials, i.e. enforcing the repeated presentation of a stimulus condition, until a minimum number of correct trials is accrued. Recording began simultaneously with the first day of training on the proper contrast discrimination task, but data analysis for the purpose of this paper was only performed from day 3 onwards, as this was the start of presenting the full range of contrasts.*

<u>Data acquisition and processing</u>. Data acquisition and preprocess details are also not critical for this study. We duplicated the texts as italics.

*Raw data were acquired at a sampling frequency of 32,556 Hz with a 24-bit analogue-to-digital converter, with minimum and maximum input ranges of 11 and 136,986 microvolts, respectively (pre-set by Neuralynx, Inc.), a DMA buffer count of 128 and a DMA buffer size of 10 ms, using a 64-channel Digital Lynx 16SX Data Acquisition System (Neuralynx, Inc.). Digital referencing of voltage signals was performed prior to the recording of raw data, using commercially provided Cheetah 5 Data Acquisition Software v. 5.4.0 (Neuralynx, Inc.), to yield good SNRs for each channel.*

*Following each recording session, the raw data were processed offline using both commercial (Neuralynx, Inc.) and custom-written (Matlab, Mathworks) software. Signals were extracted using the Cheetah 5 Data Acquisition Software. The sampling frequency remained the same (32,556 Hz), while the bandpass filter frequency and the input range settings were individually tailored to each channel. Raw data were bandpass filtered with a low cut frequency of 600 Hz and a high cut frequency of 4000 Hz and saved at 16-bit resolution. This stage of processing generated 'continuous multi-unit activity (MUA)' data, which was further processed to yield 'spiking MUA'.*

*An iterative procedure was carried out on the continuous MUA signals for each channel, in which the threshold for spike extraction was varied according to a staircase procedure, in order to yield levels of spontaneous spiking MUA (before the onset of the sample stimulus) that were similar (within 1% of a 'target' level) across sessions. To set the target level for each channel, the threshold was initially selected manually for all channels and sessions, and a 'representative' session was selected for each channel (i.e., a session with an 'average' SNR [see below for description] for that channel). Hence, the extraction of spiking MUA was performed such that spontaneous activity levels were standardized across recording sessions. As spontaneous activity levels were deliberately kept uniform across training days, we did (or could) not study whether spontaneous activity levels changed during training. What this method did allow, however, was the rigorous comparison of levels of stimulus-evoked activity across the training period, relative to spontaneous levels.*

*Receptive fields were mapped using a reverse correlation procedure 17 for each recording channel prior to training and recording. Additionally, orientation and spatial frequency tuning was determined using a reverse correlation procedure 17. RF locations and tuning preferences were highly consistent across the training period as determined by regular remapping while learning commenced (every 3–5 days).*

<u>Data exclusion</u>. We used the exactly same methods to compute SNR of individual channels and applied the same criteria to select high profile channels. The SNR computation are described as below in the original paper:

*The SNR was calculated for each channel on each day. The SNR was calculated as:*

$$SNR = \frac{Mean_{stimulus\ activity} - Mean_{spontaneous}}{SD_{spontaneous}} \quad (S3)$$

*whereby the mean stimulus activity was obtained from 150 to 250 ms after test onset, while the mean spontaneous activity was obtained during the 300-ms period before test onset. SD is the standard deviation of the mean response. This was calculated for each test contrast condition, yielding 14 SNR values per recording session for a given channel. Trials were included regardless of whether the subject' s response was correct. The size of the SNR varied depending on the test contrast. The highest of the 14 SNR values was then taken as being representative of the signal quality from a given channel for each session. Channels were included in the individual channel analyses if they had daily SNR ≥ 1, on at least 80% of the total number of recording days.*

Note that according to this criteria, 29 and 20 channels were included for monkey 1 and 2, respectively. Two channels in monkey 1 were found to show monotonically decreasing contrast response functions. We thus excluded the two channels for further analyses.

<u>Population activity analyses</u>. Previous studies have suggested that the variability of neuronal responses can be decomposed into two components—the trial-by-trial variability of firing

rate and the intrinsic Poisson variability of spike generation. We hypothesize that VPL, as a top-down task signal, mainly manifests as the changes in the trial-by-trial variability of firing rate rather than altering the intrinsic Poisson statistics of the spike because the latter component is more driven by low-level biophysical properties of neuronal spikes. We used a simple multivariate Poisson-lognormal (MPLN) model as used in previous studies 18-21, which assumes that the spike counts (SC) of individual channels are generated by a homogeneous Poisson process, given the underlying firing rate ($\overrightarrow{fr}$). The model also assumes that the underlying firing rate of channels is represented by a multivariate log-normal distribution with a mean vector $\mu$ and covariance $\Sigma$, which accounts for the covariability among channels and prevents negative firing rates. The model is expressed mathematically as:

$$SC_i \sim Possion(T \times fr_i) \tag{S4}$$

$$\overrightarrow{fr} \sim \log normal\,(\vec{\mu}, \Sigma) \tag{S5}$$

To infer the latent parameters μ and Σ, we used the variational inference method with an Adam optimizer and a learning rate of 0.1. The mean of the posterior inference via variational inference was taken as our point estimate of μ and Σ. We then defined the signal vector, signal separation, response variance, signal rotation angle, PC strength, and PC rotation angle, in a manner similar to our previous work with DCNN models and fMRI data. The only difference is that we now applied these calculations to multi-unit activity within a latent space characterized by μ and Σ. This was followed by the application of our established stepwise analysis approach.
